# Quantification and differential analysis of mass spectrometry proteomics data with probabilistic recovery of information from missing values

**DOI:** 10.1101/2025.04.28.651125

**Authors:** Mengbo Li, Simon A. Cobbold, Gordon K. Smyth

## Abstract

Mass spectrometry (MS) is the technology standard for expression proteomics, but statistical analysis of the resulting data is complicated by the occurrence of missing values. Missing values remain ubiquitous in MS-based proteomics data and are especially frequent in the emerging fields of single-cell and spatial proteomics. The limpa package implements new methods for quantification and differential expression analysis of MS proteomics data, including probabilistic information recovery from missing values. limpa summarises peptide-level data to estimate an expression value for every protein in every sample. Expression values that are supported by fewer detected peptides or involve more missing values are treated as less precise and are downweighted in the differential expression analysis, maximising statistical power while avoiding false discoveries. limpa produces a linear model object suitable for downstream analysis with the limma package, allowing complex experimental designs and other downstream tasks such as the gene ontology or pathway analysis.

## Introduction

Proteomics, the large-scale study of proteins, is one of the cornerstones of biomedical research, together with genomics, transcriptomics and metabolomics [1]. While transcriptomics has been tremendously successful for profiling gene activity, proteomics focuses more directly on the functional machinery inside and outside of cells [2]. One of the most frequently occurring and most widely useful tasks is to estimate the expression levels of all proteins in a series of biological samples and to conduct statistical tests for changes in protein expression between experimental conditions or cell types [3]. Proteomics studies have found that diverse cellular systems tend to have similar proteomes, with few proteins being uniquely detectable in specific situations [4, 5]. The identity of cellular systems is determined therefore more by the abundance at which they express their constituent proteins, rather than by the presence or absence of particular proteins, emphasizing the central role that protein differential expression analysis plays in the exploration of biological systems. Mass spectrometry (MS) has become established as the standard technology for expression proteomics, because of its accuracy, specificity and applicability across all organisms [5, 1]. Guo *et al* [1] reviewed recent improvements to MS technology across all workflow stages. Notable developments include increased use of data-independent acquisition (DIA) methods for higher coverage and precision [6, 7, 8], AI tools for peptide identification and quantification [9], and the emergence of single-cell and spatial MS-based technologies [10, 11, 12, 13, 14, 15].

Bottom-up proteomics, in which proteins are enzymatically cleaved and MS is used to identify and quantify the resulting peptides, is the most widespread proteomics workflow [16, 5, 17]. While this approach is experimentally and computationally tractable, a persistent problem that complicates downstream analyses is the fact that not very peptide is necessarily detected and quantified in every biological sample, resulting in unpredictable missing values at both the peptide and protein levels [18, 19]. While strategies such as DIA and match-by-runs have been deployed to minimise the occurrence of missing values [20], missing values remain ubiquitous in MS-based proteomics data and are especially frequent for single-cell, spatial, and blood plasma samples [21, 22, 23].

Missing values cause a dilemma for the analysis of MS expression data because they do not occur due to any easily characterizable mechanism, for example entirely at random or according to a clear detection limit [24]. This leaves researchers with the choice of either imputing the missing values and treating the imputed values on an equal footing with observed values, a practice known to lead to error rate inflation in statistics [25], or treating the missing values as random and thereby losing any information that they might contain.

From a statistical point of view, missing-data mechanisms can be classified into four general categories, from the simplest to the most complex. Missingness can be completely at random, or can depend on observed predictors, or can depend on unobserved predictors, or can depend on the missing value itself [26]. We and others have shown that the proportion of entirely random missing values is small, essentially because such a process would affect all peptides, whereas peptides at sufficiently high expression levels appear to be reliably detected with high probability [24]. Despite much literature of missing values in MS, we are not aware of any publication that has identified an observed predictor for missingness. This leaves the last two categories. In terms of unobserved predictors, it is reasonable to suppose that the probability of detection might depend to some extent on the chemical and structural properties of the protein and component peptides, and on the properties of other proteins with similar sequences and profiles. However, information relating chemical properties to detection is not currently available to the data analyst and, in the absence of such information, all the proteins and peptides must be treated equally. In any case, unobserved predictors of a structural nature should be relatively unimportant in the context of differential expression analyses, provided the predictors remain constant between experimental conditions and therefore cancel out of differential comparisons. On the other hand, it is indisputable that missingness must depend on the missing value itself, first because a peptide that is not expressed cannot be detected, but also because peptides at increasing expression levels should be increasingly easy to detect and identify. The concept of “detection limit” implies that expression values below a threshold will become missing, but we view such detection limits as stochastic. We are not aware of any strict, fixed detection limits for any MS technology. Overall, these considerations lead us to believe that, for a randomly chosen peptide, the probability that the peptide can be detected and measured, i.e., is not missing, should be a monotonic increasing function of its underlying expression value in that sample. We call this function the *detection probability curve* (DPC). This dependence has important statistical implications, because missing values therefore provide probabilistic information about the underlying expression values.

It has been observed empirically from a wide range of MS proteomics datasets that peptides at lower intensities tend to have higher proportions of missing values, confirming that missingness is intensity-dependent [24]. Such empirical relationships do not, however, define the DPC itself, which depends on expression values that are missing and hence needs to be estimated indirectly. Li & Smyth (2023) developed a strategy for DPC estimation based on the mathematical technique of exponential tilting [24]. They also showed that a logit-linear formula for the DPC appeared to be adequate, with little evidence for more complex curves. The slope of the DPC evaluates how much statistical information can or cannot be recovered from the missing value pattern, and can be used to inform downstream analyses [24].

In this article, we develop methods based on the DPC for protein quantification and for differential expression analysis of MS data. The methods are implemented in the Bioconductor software package limpa [27]. The package uses the observed proteomics data to estimate the detection probability curve (DPC), which provides a formal probabilistic model for the intensity-dependent missingness. Next, the package implements a novel protein quantification method, called DPC-quant, in which missing values are represented by the DPC. An empirical Bayes scheme is employed to borrow information across the tens of thousands of peptides measured in a typical experiment. A multi-variate normal prior is estimated empirically from data to describe the variability in log-intensities across the samples and across the peptides. Finally, quantification uncertainty is incorporated into the differential expression analysis using precision weights. Leveraging the linear modelling and empirical Bayes functionality of the limma package [28], a new variance modelling approach with multiple predictors is used, which allows the DPC-quant precisions to be propagated to the differential expression analysis while simultaneously assuming a mean-variance relationship.

Peng *et al*. [3] identified five key steps in a typical proteomics analysis: raw data quantification, expression matrix construction, matrix normalization, missing value imputation, and differential expression analysis. We further separate quantification into quantification at the peptide or precursor level and summarisation at the protein level. The limpa-limma pipeline described here starts with peptide or precursor level data from a quantification tool such as DIA-NN [9], Spectronaut [29] or MaxQuant [30], and undertakes all the other steps of the workflow. limpa first estimates the strength of the relationship between missing values and underlying intensity, which is quantified in the form of a logistic DPC. Then, protein-quantification, expression matrix construction and missing value imputation are all accomplished together as a single operation by fitting an additive model with empirical Bayes priors to the peptide or precursor level data for each protein. Importantly, this quantification step also estimates the technical precision with which each protein value is quantified, yielding a standard error that depends on the number of precursors and the percentage of missing values and the consistency of the precursors for that protein. This standard error is then propagated to the differential expression analysis, and forms the basis of a novel variance modelling step, which adds power and robustness to the differential expression analysis.

Unlike previous pipelines, limpa produces a complete protein-level expression matrix without the need for imputation as a separate step. By modelling the relationship between intensity and the missing value frequency, limpa is able to produce approximately unbiased log-expression values and log-fold-changes, even for proteins at low intensities. The resulting matrix can be used directly for downstream analyses such as clustering or network construction, although downstream methods that can take account of the quantification standard errors are preferable.

limma has been identified previously as an top-ranking differential expression tool across a range of MS technologies and imputation strategies [3]. Here we make use of limma’s ability to incorporate variance trends and precision weights derived from the missing value pattern to further refine the limma analysis pipeline, improving accuracy and increasing power to detect differential expression. The DEqMS package [31] has previously used the number of peptide-spectrum-matches (PSMs) or peptides for each protein to predict a protein-level variance trend to enhance limma’s empirical Bayes differential expression procedure. The limpa analysis presented here can be seen as a more comprehensive version of this type of approach, incorporating missing values, observation-specific precision weights and a more graduated assessment of quantification uncertainty.

The limpa package is fully compatible with limma pipelines, allowing any arbitrarily complex experimental design and other downstream tasks such as the gene ontology or pathway analysis. By leveraging the linear modelling functionality of the limma package, the limpa-limma pipeline includes integrated access to a wide range of advanced analysis tools including arbitrarily complex experimental designs, correlated samples, sample quality weights and gene set enrichment. The pipeline allows for independent groups, paired comparisons, batch effects or continuous covariates, in an integrated fashion.

To our knowledge, there have been three previous attempts to incorporate DPCs into proteomics differential expression procedures. Luo *et al*. [32, 33] assumed a logistic DPC as part of a Bayesian model for differential expression analysis of iTRAQ data. O’Brien *et al*. [34] assumed a probit DPC as part of a Bayesian model for differential expression testing. Ahlmann-Eltze and Anders [35] assumed sample-specific probit DPCs as part of an empirical Bayes model for differential expression testing. In all cases, the DPCs were treated a nuisance parameter in a Bayesian testing procedure and were not examined or interpreted in their own right. The Luo method is specifically for iTRAQ while the second and third work on protein-level summaries and do not produce quantifications. The methods of Luo and O’Brien are not implemented in publicly available software as far as we know.

## Results

### The DPC summarises known and unknown mechanisms

The starting point for a limpa analysis is a file containing peptide intensities from a series of biological samples, generated by a quantification tool such as DIA-NN [9], Spectronaut [29] or MaxQuant [30]. Almost always, there are peptides that are detected and quantified for some samples but not others, leading to missing values when the intensity data is summarised in matrix form with rows corresponding to peptides and columns to samples. The observed intensities can be viewed as incomplete data [36, 37]. We imagine conceptually that there is a complete data matrix that could have been measured had the MS technology been “perfect”, without any detection limits, and the observed data arises as a randomly censored version of the complete data. The probability that a complete data intensity is non-missing in the observed data is assumed to be a monotonic increasing function of the complete data value itself. While the exact form of this function is not important, for simplicity and mathematical tractability we assume that this function, called the DPC, has a logistic function form. The probability that peptide *p* in sample *i* is detected is assumed to be *F* (*β*_0_ + *β*_1_*y*_*pi*_), where *F* is the logistic function, *y*_*pi*_ is the complete data intensity and *β*_0_ and *β*_1_ are intercept and slope parameters that are specific to each dataset, depending on the MS technology used [24].

The higher the slope *β*_1_ of the DPC, the more information about the complete intensity can be inferred from the fact that it is missing [24]. A zero slope (*β*_1_ = 0) represents missing at random, in which case the missing values can simply be ignored without loss of information. A very large positive slope, together with a large negative intercept, would correspond to left-censoring with a fixed detection limit. More generally, the DPC can be viewed as arising from variable detection limits, whereby each peptide has its own detection limit, but the limits are unknown and vary between peptides and samples according to a logistic distribution. In this interpretation, the slope of the resulting DPC is inversely related to the variance of the detection limit distribution.

In general, the DPC does not assume any particular physical mechanism causing the missingness, but represents the aggregate probabilistic effect of all the factors that might be at play.

### The complete normal and observed normal models

Li & Smyth (2023) assumed a normal distribution model for the observed log-intensities between replicate samples, an assumption that underlies many common analysis strategies. In the current article, we assume instead that the complete data log-intensities are normally distributed, an assumption that seems natural and is more mathematically consistent for the purposes of quantification and differential expression. We call these two approaches the “observed normal” (ON) and “complete normal” (CN) models, respectively. If we consider a set of independent replicate samples under the same experimental conditions, the CN model implies that the observed log-intensities for any given peptide will follow a slightly skew distribution, because the samples with low intensities in the complete data tend to be missing in the observed data. On the other hand, the ON model implies that the complete data distribution is a mixture of two normal distributions, with equal variances but shifted means [24]. While these results show that the two models are mathematically different, the quantitative differences between the two models are small from a practical point of view, so much so that they can’t be distinguished from real data (Fig. 1).

**Figure 1:**
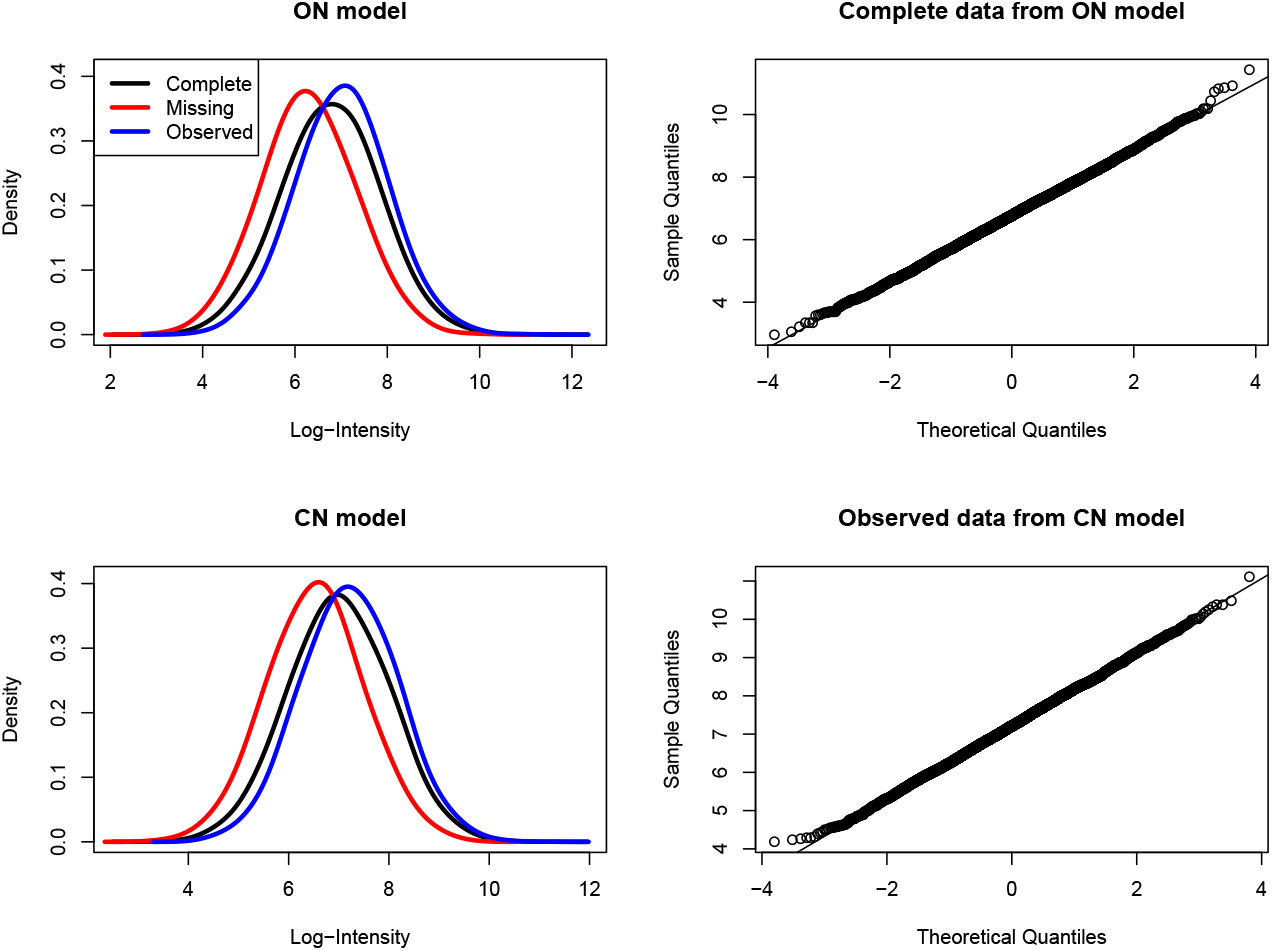
Observed-normal and complete-normal models give similar results. The top two panels show results from an ON model where the observed values are normally distributed with mean 7 and standard deviation 1. The normal probability plot shows that the complete values are also normally distributed to a close approximation. The bottom two panels show results from a CN model where the complete values are normally distributed with mean 7 and standard deviation 1. A normal probability plot shows that the observed values are also normally distributed to a close approximation. Density plots show that the observed and missing value distributions have similar standard deviations but with shifted means. Results are from 10,000 simulated values for each model with DPC intercept and slope equal to −4 and 0.7 respectively. The percentage of missing values is 34% for the ON model and 31% for the CN model.

### Quantifying protein expression

limpa uses the DPC, together with a Bayesian model, to estimate the expression level of each protein in each sample. It fits an additive model

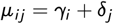

to the peptide log-intensities for each protein, where *µ*_*ij*_ is the expected log-expression of peptide *j* in sample *i, γ*_*i*_ is the log-expression level of the protein in sample *i* and *δ*_*j*_ is the baseline effect for peptide *j*. A sum-to-zero constraint is applied to the peptide effects *δ*_*j*_ so that the protein expression *γ*_*i*_ represents the average log-expression of the peptides in sample *i*. The log-likelihood consists of squared residuals for each observed peptide value and the log probability of being missing for each non-detected peptide value. A multivariate normal prior is also applied to the protein log-expression values, where the prior is estimated from the global data for all proteins. The prior distribution assumes that average log-intensity is normally distributed across peptides, and that log-intensities for the same peptide are positively correlated. DPC-Quant maximises the log-posterior with respect to the *γ*_*i*_s and *δ*_*j*_s, and the final *γ*_*i*_s become the protein quantifications. DPC-Quant also returns the posterior standard error with which each log-expression value is estimated.

### DPC-Quant gives reliable log fold change estimates

Using the mixed-species dataset, we assessed the accuracy of protein quantification by comparing the log fold change (logFC) estimate of each protein to its theoretical value. According to the mixing ratios of three species in each group (Fig. 2A), the theoretical logFC(B/A) values are 0, log2(30/3) and log2(3/30) for human, *E. coli* and yeast respectively. Therefore, we expect all *E. coli* proteins to be up-regulated and all yeast proteins to be down-regulated comparing group B to group A.

**Figure 2:**
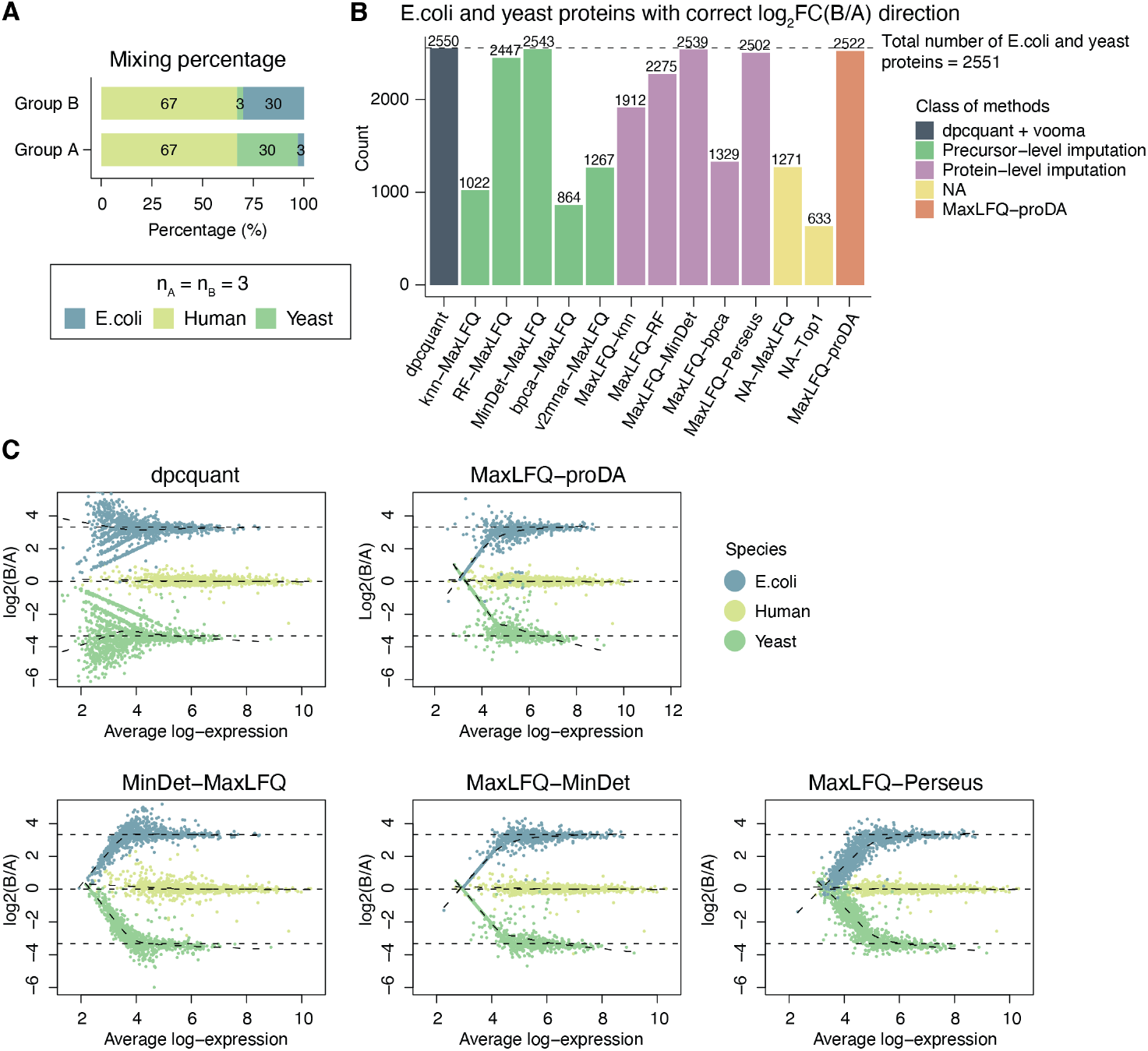
Quantification accuracy evaluated on the mixed-species dataset. (A) Experimental design. (B) Numbers of *E. coli* and yeast proteins with the correct log2FC direction. (C) Mean-difference (MD) plots for the five best-performing methods in panel B. Complete results are shown in Supplementary Fig. 4.

Fig. 2B summarises the number of yeast and *E. coli* proteins with the correct logFC direction by each pipeline. DPC-Quant outperformed all existing pipelines, giving the correct logFC sign for 2,550 out of the total of 2,551 *E. coli* and yeast proteins that were quantified. MinDet was the next best, with comparable performances applied on both precursor and protein levels. The two NA-approaches were among the least performing methods, mostly because many proteins were completely missing in one of the groups. As a result, logFCs were not estimable for such proteins. Among the proteomics specialist methods, MaxLFQ-Perseus and MaxLFQ-proDA had comparable results and considerably outperformed v2mnar-MaxLFQ. Among the generic imputation methods, knn performed markedly better when applied on the protein level, while bpca had the poorest performance on both levels. RF and MinDet showed similar performances applied on precursor- and protein-level data.

We next compared each logFC estimate to its theoretical value using the mean-difference (MD) plots. The three dashed lines in each MD plot (Fig. 2C), from top to bottom, indicate the expected logFCs for *E. coli*, human and yeast proteins respectively. For each species, we also fitted a lowess curve for the logFC estimates against the average log-intensities. If the logFC estimates are accurate, the lowess fit should be approximately horizontal at the corresponding theoretical logFC value.

DPC-Quant gave the most truthful logFC estimates for all species, with each lowess curve closely tracing the corresponding theoretical value (Fig. 2C, Supplementary Fig. 4). All pipelines gave accurate logFC estimates for the human proteins, which have less missingness than the other species at the precursor-level data. Both MinDet-based pipelines gave systematically biased logFC estimates for *E. coli* and yeast proteins at low intensities. The two proteomics specialist pipelines, MaxLFQ-proDA and MaxLFQ-Perseus, also gave biased logFC estimates for the *E. coli* and yeast proteins at low intensities. Both NA-pipelines were able to reliably estimate the logFC for *E. coli* and yeast proteins, although the estimates were slightly biased toward zero in low-abundance proteins (Supplementary Fig. 4). Note that although the same number of protein IDs were retained for all pipelines but v2mnar-MaxLFQ (Supplementary Fig. 3E), a lot of proteins were missing from the MD plots for the two NA-pipelines. The missing proteins were completely missing in one of the groups for which the logFCs were not estimable. Meanwhile, less proteins IDs were retained by v2mnar-MaxLFQ, because v2mnar by default only includes precursors that are observed in more than 4 samples.

In addition, the accuracy in logFC estimates also depends on average log-intensity. To illustrate this, proteins were binned by their average log-intensities, and the logFC distribution was visualised within each bin. As shown in Supplementary Fig. 5, all pipelines gave consistently accurate logFC estimates for human proteins at all abundance levels. However, for *E. coli* and yeast proteins, DPC-Quant was the only pipeline that consistently yielded accurate logFC estimates at all intensity levels. All other pipelines were only able to give the correct logFC estimates for proteins of higher abundances.

### Propagating uncertainty to the differential expression analysis

When summarising precursors into the protein level, different precursors may be missing in different samples; and different numbers of missing precursors may occur to different samples. Meanwhile, similar to other omics technologies, the variance of log-intensity values also depends on the protein expression level. For these reasons, protein intensities can vary greatly from sample to sample within each protein and as the result, the variances of individual summaries are unequal. Therefore, we apply precision weights at the observational level to downweight unreliable protein intensities in DE analysis. In order to choose these weights, we need to model the variance in protein intensities.

Similar to microarray and RNA-seq data, the variance in log-intensities tends to be greater in proteins of low average expression. As pointed out in Law *et al*. [38], it is key to correctly model the mean-variance relationship in DE analysis to maintain FDR control while improving the statistical power. We observe a similar inverse relationship between the square root of residual standard deviation and the average log-intensity in each protein in proteomics data, regardless of the summarisation algorithm used for protein quantification (Supplementary Fig. 1A).

Bottom-up MS data needs to be summarised at the protein level. Protein-level intensities are obtained for each protein in every sample by summarising their precursor-level log-intensities, where missing values lead to quantification uncertainty. The variance of protein intensities depends strongly on quantification uncertainty, which depends on the number of precursors and on the level of missingness at the precursor level. The protein-level summary of samples with more missing values in the precursors are less reliable and tend therefore to have greater variances. In DPC-Quant, we measure quantification uncertainty by the standard error associated with each protein-level estimate, which is output alongside the protein-level summary by our software. Protein intensities of greater quantification uncertainty have larger standard errors. Strong uncertainty-variance trends can be seen in real data as shown in Supplementary Fig. 1B.

We use a bivariate linear model to summarise how the variance in protein intensities depends jointly on average expression and the quantification uncertainty. As explained in Law *et al*. (2014) [38], a technical limitation exists where although we wish to estimate the variance of each individual observation, it is fundamentally impossible to have replications for each single observation so that the variance can be empirically estimated. Thereby to obtain observational weights, we first estimate the trend in residual variances on the protein level. The fitted trend is then interpolated to predict the variance of each individual observation, which is used as an inverse weight for the corresponding protein quantity. This first step is called variance modelling and the second step is weight interpolation, both of which are implemented in the limpa function voomaLmFitWithImputation(). Supplementary Fig. 1C demonstrates the vooma trend in the human mixture dataset.

### limpa controls type I error rate correctly

Using a mixture design, limpa was then evaluated and compared to several widely adopted DE pipelines. In the human mixture dataset, total proteins derived from normal bladder, frontal lobe and cerebellum tissues were mixed at 6 different proportions (Fig. 3A). The bladder tissue was used as the constant background as we varied the cerebellum to frontal lobe ratio for each condition group. Four technical replicates were generated at each mixing ratio. There were 15.8% missing values on the precursor level after filtering out the non-proteotypic precursors and precursors mapped to the compound protein groups. The estimated DPC coefficients were *β*_0_ = 8.00 and *β*_1_ = 0.74 (Supplementary Fig. 6C).

**Figure 3:**
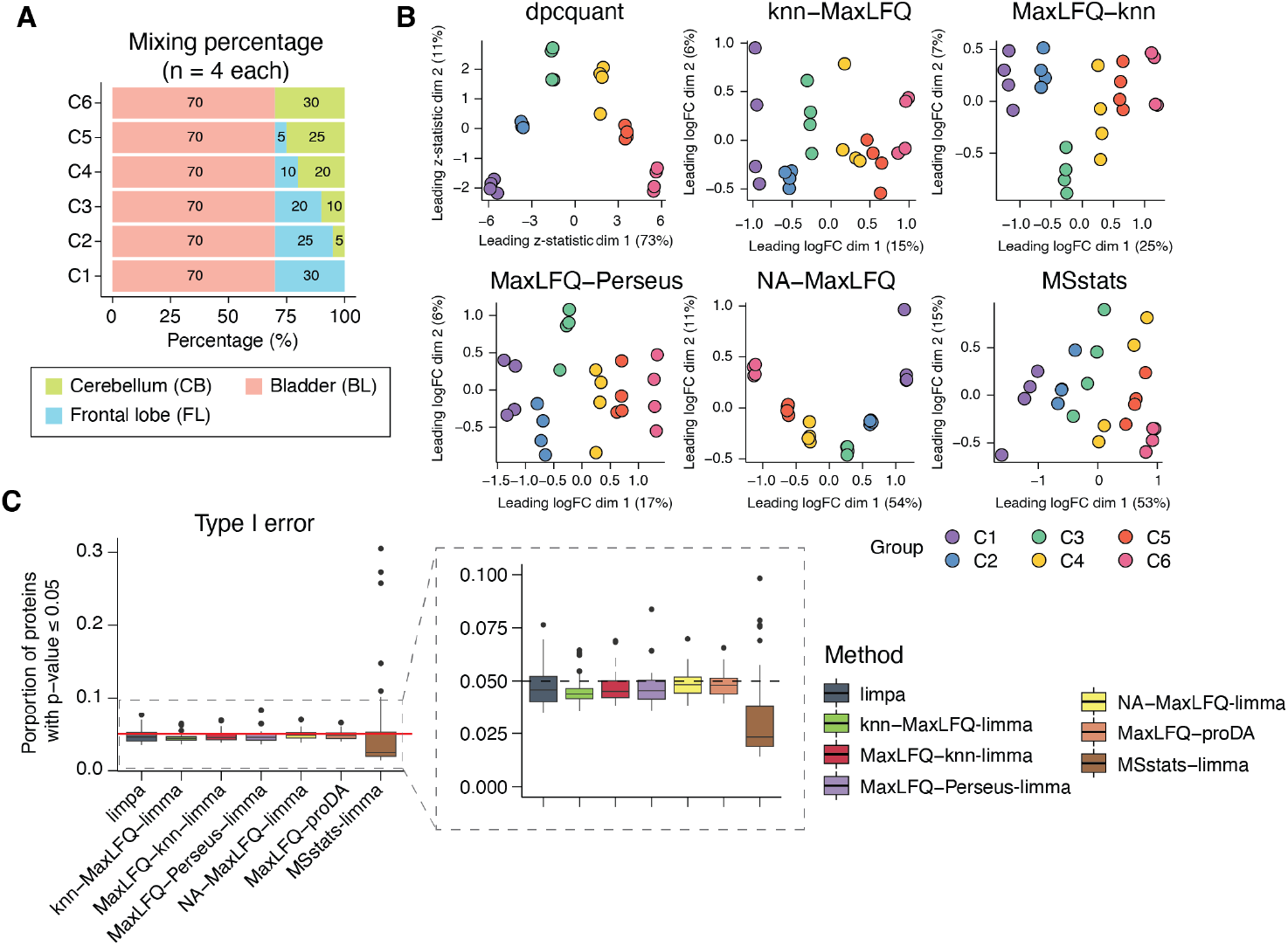
Protein quantification and DE analysis pipelines evaluated on the human mixture dataset. (A) Experimental design. (B) Multidimensional scaling (MDS) plot on protein quantification by each method. (C) Type I error rates over 50 simulated null comparisons at the p-value cutoff of 0.05.

Protein summarisation was performed to obtain the protein-level data. The MDS plot on the protein quantification by each pipeline is shown in Fig. 3B. Condition groups were separated mainly along the x-axis for all pipelines. DPC-Quant had the highest percentage of variance explained by the first dimension compared to all other pipelines. Among the rest of the pipelines, NA-MaxLFQ had the least technical variation within in each condition. However, there was one obvious outlying sample in group 1.

Error rate control was evaluated based on null comparisons where no genuine DE should exist. Replicates of each condition were randomly split into two subgroups over 50 simulations. DE analysis was performed where an additive model with two covariates, condition and subgroup, was fitted for each protein. No DE should exist between subgroups of the same condition, as they are technical replicates. For the subgroup coefficient, the proportion of nominal p-values that was less than or equal to 0.05 was calculated to estimate the type I error rate of each simulation. The default MSstats pipeline was not included in this evaluation because it does not accommodate additive models. Fortunately, MSstats returns the protein-level data, where proteins are summarised using Tukey median polishing and missing values are imputed using an accelerated failure time model. Therefore, null comparisons could be performed on the protein-level summarisation by MSstats using limma (MSstats-limma).

The distribution of type I error rates from 50 simulations is visualised in Fig. 3C. Overall, all pipelines controlled type I error rate correctly, although MSstats-limma had more variable results across simulations. In most simulations, MSstats-limma was more conservative compared to the rest of the pipelines, but there were also instances where it became overly liberal. The average and median type I error rates over 50 simulations were also calculated at different nominal p-value cutoffs (Supplementary Fig. 6D,E). Results were qualitatively similar at various p-value cutoffs.

### limpa has the best power and the lowest false discovery rate

Next, we assessed the power and the false discovery rate (FDR) to detect DE proteins of each pipeline. DE analysis was performed to compare groups C6 versus C1, and DE proteins were defined by the FDR cutoff for significance at 0.05. The number of DE proteins identified by each pipeline was summarised in Supplementary Fig. 6G. The number of protein IDs quantified by each pipeline and the number of protein IDs with a testable logFC between C6 and C1 were summarised in Supplementary Fig. 6F.

True and false discoveries were defined based on the proteomic profiling of the pure tissue lysates (the pure tissue dataset). Five technical replicates from the total proteins derived from normal bladder, frontal lobe and cerebellum tissues were analysed (Supplementary Fig. 7). Protein quantification was performed by each pipeline, following which DE analysis was performed to compare the pure cerebellum samples to those of frontal lobe. DE proteins were defined at 5% FDR, which were assumed to be the true DE proteins when comparing C6 versus C1 in the human mixture dataset. The number of true DE proteins corresponding to each pipeline was summarised in Supplementary Fig. 7G. Meanwhile, proteins that were not significant were assumed to be the true non-DE proteins.

The power to detect to true differential expression was evaluated by ranking the proteins by their p-values. Pipelines were compared in terms of the number of true discoveries at each given number of proteins selected as DE. Good performing methods should rank the true DE proteins ahead of the non-DE proteins. All pipelines had similar performances until a protein rank of around 2,000, but limpa discernibly outperformed all other pipelines at greater cutoffs (Fig. 4A). Such an observation was not surprising because this human mixture dataset did not have a lot missing values, whereas most pipelines included in this evaluation were only different in how they handle missing values. In fact, when no imputation was performed, there was only 8% missingness on the protein level by MaxLFQ summarisation (NA-MaxLFQ).

**Figure 4:**
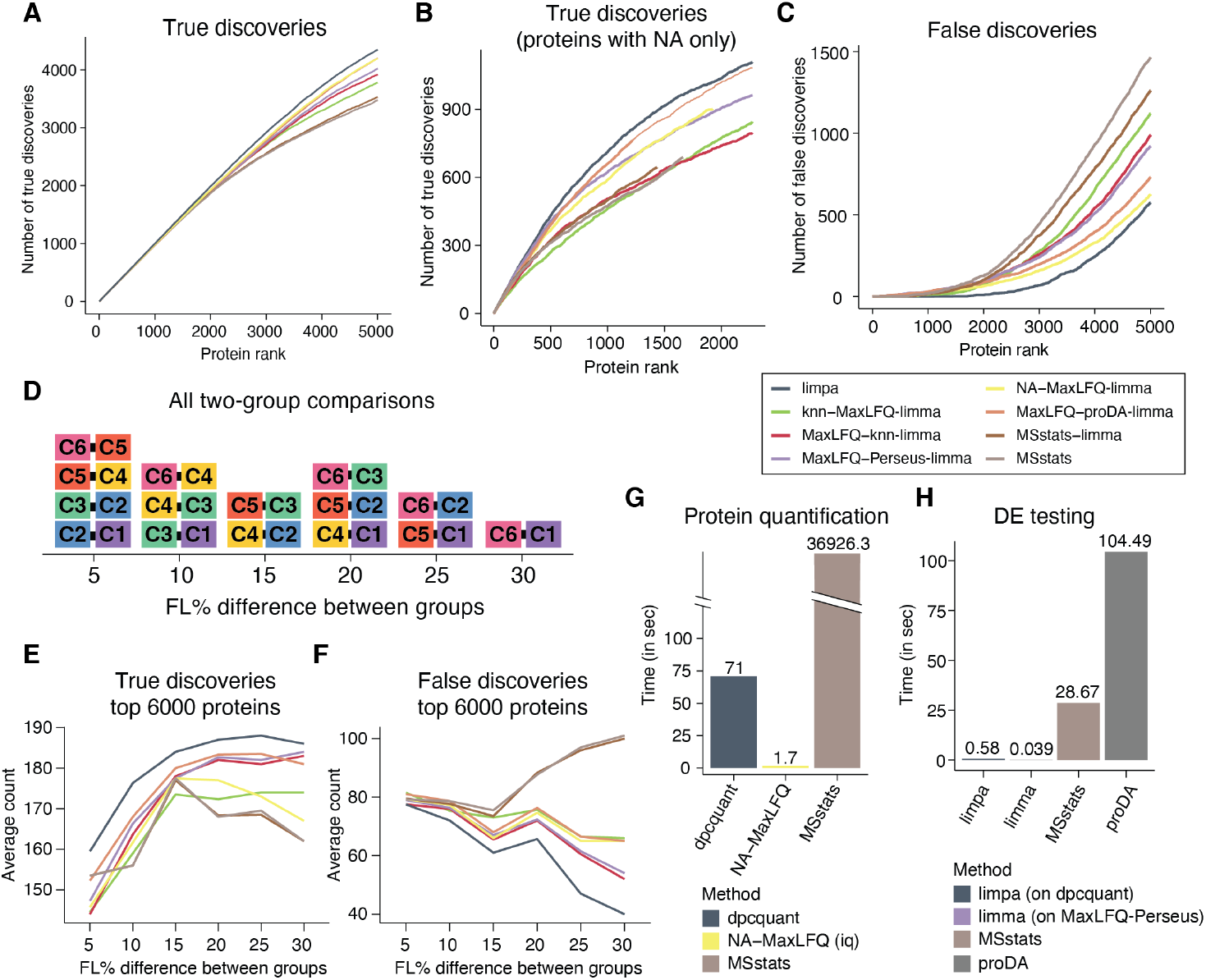
DE analysis pipelines evaluated using the human mixture dataset. (A) True discoveries. The number of true discoveries is plotted for each method versus the number of proteins selected as DE. (B) True discoveries. The number of true discoveries is plotted for each method versus the number of proteins selected as DE, only considering proteins with any missingness if quantified by NAMaxLFQ. (C) False discoveries. The number of false discoveries is plotted for each method versus the number of proteins selected as DE. (D) Summary of all possible two-group comparisons in the mixture design. The comparisons are categorised by the difference in mixing percentages of frontal lobe lysates between groups. (E) Consistency of true discoveries. Proteins are ranked by their p-values. True discoveries are defined as the frontal lobe- and cerebellum-specific proteins according to the missingness patterns in the paired pure tissue lysate dataset (Supplementary Fig. 7H). The average count of true discoveries in top 6,000 proteins is calculated at each level of heterogeneity between groups in the two-group comparisons. (F) False discoveries. Proteins are ranked by their p-values. False discoveries are defined as the bladder-specific proteins according to the missingness patterns in the paired pure tissue lysate dataset (Supplementary Fig. 7H). The average count of false discoveries in top 6,000 proteins is calculated at each level of heterogeneity between groups in the two-group comparisons. (G) Computation time in seconds spent on the protein quantification step by DPC-Quant, NA-MaxLFQ and MSstats. (H) Computation time in seconds spent on DE testing by limpa, limma-trend (on protein-level data summarised by the MaxLFQ-Perseus pipeline for this evaluation), MSstats and proDA.

In order to evaluate the performance of each pipeline on proteins susceptible to missing values, we then considered the subset of true discoveries that had missing values. Specifically, we filtered for true discovery proteins that had at least 1 missing value in the NA-MaxLFQ summarisation. It then became apparent that limpa had the best power in detecting true DE proteins when missing values are a concern (Fig. 4B).

FDR was also evaluated from the protein ranking point of view. The number of false discoveries at each number of proteins selected as DE was visualised in Fig. 4C. Results showed that limpa gives the best protein ranking, with the lowest FDR for any given number of proteins selected.

### limpa gives consistent DE results on the mixture design

The mixture design also allows us to evaluate the consistency of DE results as the two mixtures of each two-group comparison become more heterogenous. For each pairwise comparison, we can calculate the difference in the mixing percentages in terms of frontal lobe as a measurement of heterogeneity between groups (Fig. 4D). DE analysis was performed for all possible pairwise comparisons by each pipeline.

True and false discoveries were defined leveraging the pure tissue lysate experiment. We assumed tissue-specific proteins should be mostly observed in the technical replicates of the tissue of interest and mostly missing in other tissue samples. Tissue-specific proteins can serve as a consistent set of true and false discoveries for evaluation of all different pipelines. The number of tissue-specific proteins are summarised in Supplementary Fig. 7H. According to our mixture design, bladder-specific proteins are the false discoveries; cerebellum- and frontal lobe-specific proteins are the true discoveries in all two-group comparisons.

The average numbers of true (Fig. 4E) and false (Fig. 4F) discoveries in the top 6,000 proteins ranked by p-values were calculated at each level of heterogeneity. We saw that limpa had the best power and the lowest FDR at all levels of heterogeneity.

### limpa is computationally efficient

We also evaluated the computation time for both protein quantification and DE testing using the human mixture dataset. All methods were timed using their default settings. DPC-Quant was completely implemented in R, and it took approximately 71 seconds to quantify 108,197 precursors into 8,230 proteins in 24 samples. MaxLFQ implemented by the iq R package was the fastest while MSstats was the slowest, although MSstats can be sped up if multiple cores are available. Another reason why MSstats was more computationally demanding is because MSstats imputes and summarises the fragments, rather than the precursors, into proteins. Although not the fastest, DPC-Quant was computationally efficient for local analyses on a laptop.

The computational time on DE testing was evaluated by the two-group comparison between C6 and C1, where there were 4 replicates in each group. The number of proteins included in the DE analysis was 8,230 for limpa, limma (on MaxLFQ-Perseus) and proDA, and 7,559 for MSstats (Supplementary Fig. 6F). The limpa and limma pipelines were easily the fastest methods for DE testing, followed by MSstats. The proDA pipeline was the slowest which took around 104 seconds.

### Cell cycle proteomes by single cell proteomics

Next we compared limpa with the popular MaxLFQ-Perseus-limma pipeline using a public single cell proteomics dataset. Populations of HeLa cells enriched at specific cell cycle stages (G1, G1S, G2 and G2M) were produced via drug perturbation [11]. Single cell proteomes were measured by the true single-cell-derived proteomics (T-SCP) workflow, and raw MS data were processed by DIA-NN [11]. After pre-processing, approximately 51.2% of data were missing on the precursor level. Fig. 5A shows the distribution of the proportion of missing values in each cell on the precursor level. Cells in G1 had the highest level of missingness compared to the rest of the stages. G1S cells had the lowest amount of missing values across all groups, yet there were still more than 25% precursors missing for most G1S cells.

**Figure 5:**
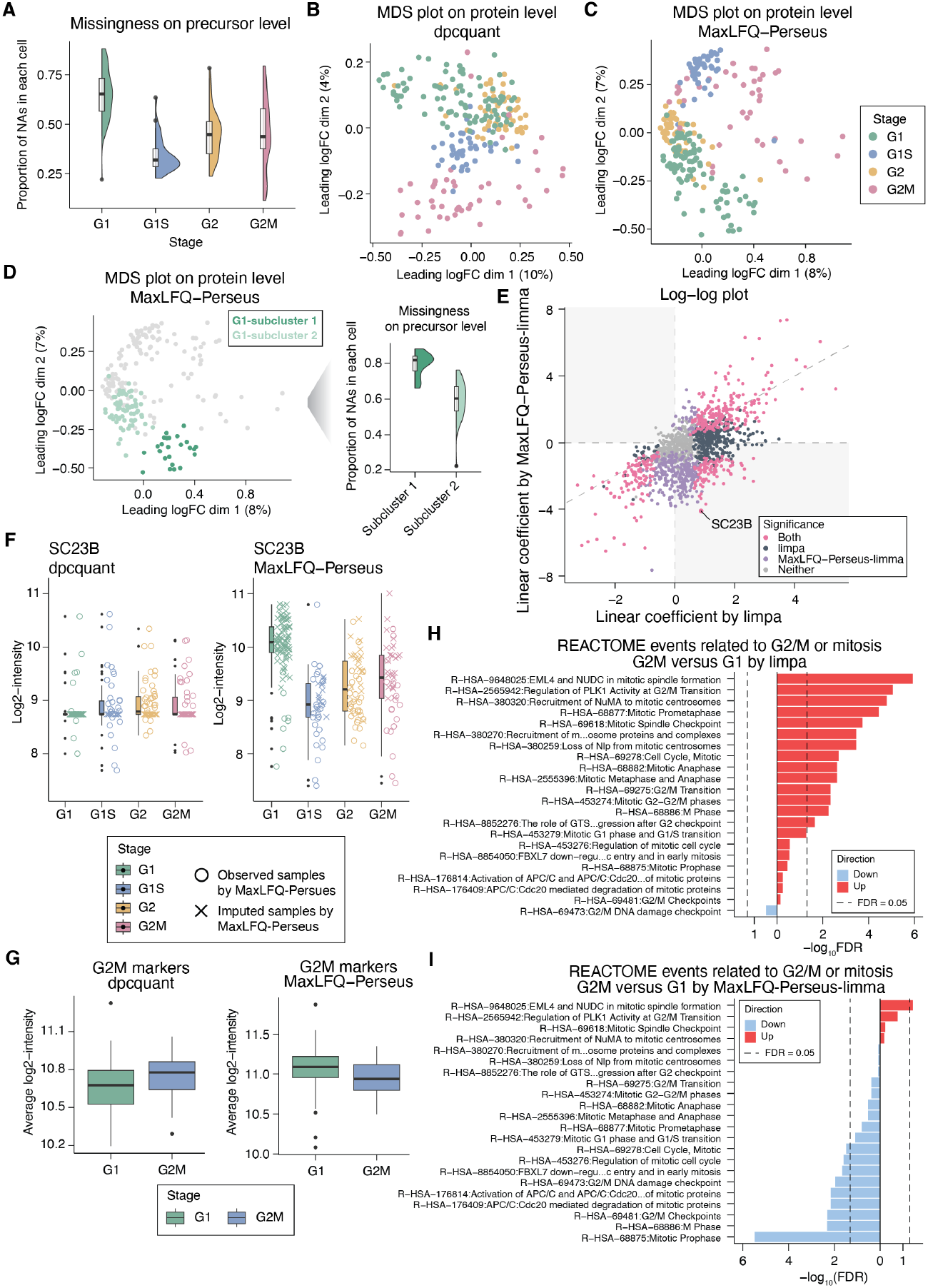
Single cell proteomic profiling of the cell cycling stages. (A) Distribution of the missing value proportion in each cell on the precursor level. (B) MDS plot on the protein level data summarised by DPC-Quant. (C) MDS plot on the protein level data summarised by the MaxLFQ-Perseus pipeline. (D) MDS plot on the protein level data summarised by the MaxLFQ-Perseus pipeline with cells from G1 highlighted (left). Distribution of the missing value proportions in G1 cells on the precursor level (right). (E) Log-log plot on the linear term. (F) Protein-level summary of each cell for the SC23B protein by DPC-Quant (left) and MaxLFQ-Perseus (right). (G) Average log2-expression of the G2M stage markers in G1 and G2M cells by DPC-Quant(left) and MaxLFQ-Perseus (right). (H) FRY gene set test on REACTOME events related to G2/M or mitosis by on protein-level data summarised by DPC-Quant. (I) FRY gene set test on REACTOME events related to G2/M or mitosis by on protein-level data summarised by MaxLFQ-Perseus.

MDS plots were generated on protein-level data summarised by both DPC-Quant and MaxLFQ-Perseus. Based on protein summarisation by DPC-Quant (Fig. 5B), while G2 cells had the greatest within-group variation, clusters of cells at each cycling stage were reasonably well-separated from each other. For protein quantification by MaxLFQ-Perseus (Fig. 5C), cell clusters were mostly separated by the second dimension with a few obvious outliers of G2M cells. In particular, there were two clear subclusters within the G1 group (Fig. 5D). We found that G1 cells in subcluster 1 had much more missing values on the precursor level than those in subcluster 2, so that the imputed values by Perseus were driving the separation between two subclusters.

We then explored the proteomic changes throughout different cell cycling stages. To do that, we assigned a numeric code for each group, that is, 1 for G1, 2 for G1, 3 for G2 and 4 for G2M, An orthogonal polynomial of degree 2 was generated based on the numeric coding of the cell cycle stages, which was then used as the design matrix. DE analysis was performed to identify proteins that significantly change across different cell cycle stages, using limpa and limma for MaxLFQ-Perseus summarisation. Fig. 5E shows the log-log plot for the linear term of the polynomial. We saw that a lot of DE proteins identified by both pipelines had opposite directions in their linear coefficients estimated by the two pipelines. For instance, according to MaxLFQ-Perseus quantification, SC23B expression markedly decreased throughout G1, G1S, G2 and G2M, while based on limpa, its expression only slightly increased with a much smaller logFC. Fig. 5F visualises the protein summary of SC23B by both methods, where the crosses indicate cells that had a missing value by NA-MaxLFQ. It was found that the large negative coefficient according to MaxLFQ-Perseus was primarily influenced by the imputed values in G1. Supplementary Fig. 9 gave more proteins similar to the SC23B example, where the imputed values in G1 cells led to biased estimates of the linear coefficient.

Next we performed a two-group comparison between G1 and G2M cells. To start with, we investigated the expression of G2M markers in this single cell proteomics dataset. G2M markers were obtained from the Seurat package [39, 40], and Fig. 5G displays the average log2-intensities of G2M markers in G1 and G2M cells by both methods. G2M markers showed higher average expression in G2M compared to G1 based on DPC-Quant as per the expectation. However, we found that G2M markers had lower average expression in G2M than in G1 based on the protein summarisation by MaxLFQ-Perseus.

We also performed FRY gene set tests on a list of REACTOME events that are related to G2/M or mitosis (Fig. 5H, I). Such a list was curated by searching for keywords including G2/M, mitosis, mitotic or the M phase. At 5% FDR, all of the significant REACTOME events were up-regulated based on limpa analysis, whereas all but one significant event were down-regulated using the MaxLFQ-Perseus-limma pipeline. In particular, the top 1 significantly down-regulated REACTOME event identified by MaxLFQ-Perseus-limma was mitotic prophase, during which chromatin condenses, centromere separates and mitotic spindle begins to form. Such processes are specific to mitotic progression, and the pathway is expected to be up-regulated in G2/M compared to G1. However, the opposite was reported based on the MaxLFQ-Perseus quantification. Additionally, the mitotic prophase event is a sub-event of the M Phase pathway, which was also reported to be significantly down-regulated by MaxLFQ-Perseus-limma. Meanwhile in the limpa analysis, both mitotic prophase and M phase events were up-regulated, and the M phase event was significant at 5% FDR (FDR = 0.006).

### Immune cell profiling in cutaneous drug reactions by spatial proteomics

Next we analysed the spatial proteomics data of immune cells in cutaneous adverse drug reactions (CADRs) from Nordmann *et al*. (2024) [15]. Deep visual proteomics (DVP) [41] was performed on FFPE lesional skin biopsies from patients with maculopapular rash (MPR), drug reaction with eosinophilia and systemic symptoms (DRESS), toxic epidermal necrolysis (TEN) and healthy individuals [15]. CD45^+^ immune cells were identified using immunofluorescence staining, following which individual cells were captured by AI-guided laser microdissection [15]. Cells from each patient were combined and analysed by LC–MS/MS [15]. The study was carefully designed that for a lot of patients, duplicates were obtained from the same patient whenever possible.

In this case study, we focus on delineating the proteomic differences between MPR versus TEN. In the original analysis, Nordmann *et al*. (2024) [15] had combined duplicates by taking the average profile for each patient. No DE protein was detected when comparing the averaged samples between MPR and TEN [15]. Hereby we demonstrate how limpa unlocks the limma suite of methods and improves the statistical power of DE analysis in real-world examples. We present an alternative analysis where all replicates were kept as is, and the correlation between biological replicates were estimated by fitting a mixed linear model for each gene [42]. A consensus correlation for all genes is calculated and incorporated into variance modelling, which greatly improves the precision of DE tests [42].

The MDS plot on DPC-Quant summarisation is illustrated in Fig. 6A, where samples are coloured by disease groups. Groups were mostly clustered along the second dimension, with a clear separation between healthy and the CADRs groups. There were a few obvious outlying samples from each group along the first dimension, and we were interested in understanding why they were outliers. We performed a weighted surrogate variable analysis, where surrogate variables that explain a high proportion of the residual variability for many of the genes are calculated [43]. We plotted the MDS plot on DPC-Quant summarisation again in Fig. 6B, and samples are now coloured by the first surrogate variable (SV1). We see that along the first dimension, SV1 values decreased. There was a clear separation by SV1 between the outliers and the rest of the samples, where the outliers corresponded to higher SV1 values.

**Figure 6:**
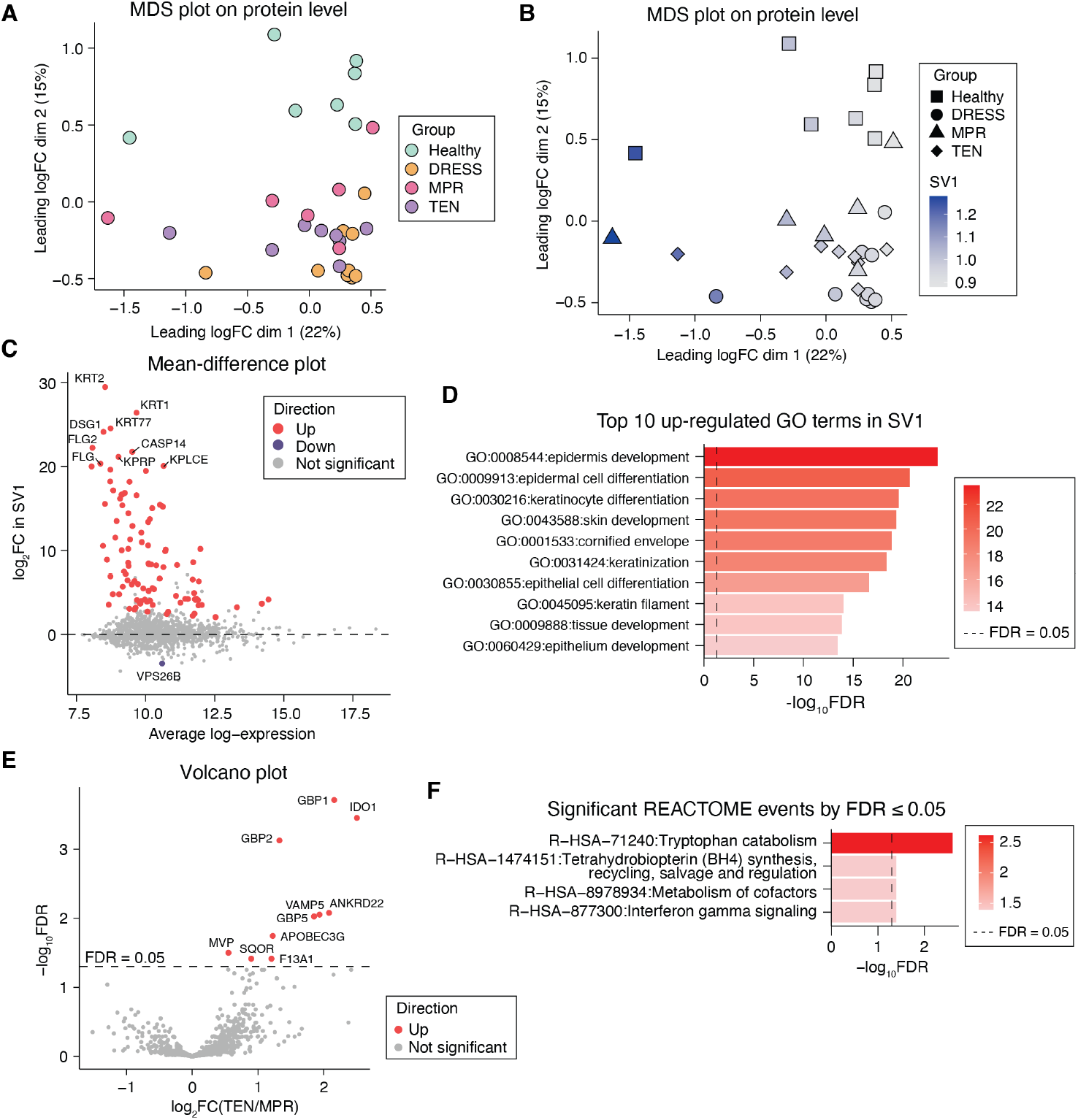
limpa analysis of spatial proteomic profiling of immune cells in healthy, DRESS, MPR and TEN. (A) MDS plot on protein summarisation by DPC-Quant. Samples are coloured by condition groups. (B) MDS plot on protein summarisation by DPC-Quant. Samples are coloured by the surrogate variable (SV) value. (C) Mean-difference plot for the surrogate variable. (D) Top 10 up-regulated Gene Ontology (GO) terms associated with the surrogate variable. (E) Volcano plot for DE proteins between TEN and MPR. (F) Top 20 up-regulated Gene Ontology (GO) terms in TEN compared to MPR.

A DE analysis was performed to identify proteins whose expression levels significantly changed with respect to the surrogate variable using limpa. The limpa analysis identified 102 up-regulated proteins and 1 down-regulated protein for SV1 at 5% FDR (Fig. 6C). Keratin 2 (KRT2), keratin 1 (KRT1) and keratin 77 (KRT77) were among the DE proteins with the largest logFCs, which are expressed in epidermal tissues. We further investigated the DE proteins associated with SV1 by performing a gene ontology (GO) analysis, and Fig. 6D displays the top 10 up-regulated GO terms in SV1. It was found that many of these GO terms were related to the biological processes involving the epidermal tissue and keratinocytes, such as epidermis development (FDR = 3.2 × 10^−24^) and keratinocyte differentiation (FDR = 2.5 × 10^−20^). Given these immune cells were extracted from the dermo-epidermal junction [15], these outliers were likely to be obtained from regions closer to the epidermal tissues. Therefore, the surrogate variable must be taken into account when comparing different diseases groups.

Following that, we performed the DE analysis to compare the CD45^+^ immune cells from TEN versus MPR patients. Both are CADRs conditions but MPR is milder compared to TEN [15]. limpa identified 10 DE proteins, and all are up-regulated in TEN compared to MPR (Fig. 6E). The top 1 DE proteins was guanylate-binding protein 1 (GBP1), which plays a key role in regulating cytokine responses. In addition, GBP1 was also reported to be significantly up-regulated in TEN compared to the healthy controls in Nordmann *et al*. (2024) [15]. The second most significant protein was indoleamine 2,3-dioxygenase 1 (IDO1), which is an enzyme that depletes tryptophan and produces immunosuppressive metabolites [44]. In particular, guanylate-binding protein 5 (GBP5) was also significantly up-regulated in TEN compared to MPR in our limpa analysis, whereas GBP5 was detected only in TEN patients therefore was not testable in the original analysis [15].

Fig. 6F illustrates the significantly differentially regulated REACTOME pathways in TEN compared to MPR at 5% FDR. The top 1 pathway was tryptophan catabolism, which plays a significant role in MPR through macrophages. It was reported in Nordmann *et al*. (2024) [15], IDO1, a key regulator in tryptophan catabolism, had the higher expression in macrophages in TEN compared to the healthy controls. Additionally, Nordmann *et al*. (2024) [15] also showed that compared to the healthy, macrophages in TEN had higher expression of proteins related to the interferon pathway, including IDO1 and GBP5. Both proteins were also significantly up-regulated in the limpa analysis comparing TEN to MPR (Fig. 6E). Nordmann *et al*. (2024) [15] suggested that this potentially reflected the higher expression of interferon gamma receptor. In fact, the interferon gamma signalling pathway was identified as significantly up-regulated in TEN compared to MPR in the limpa analysis. Further experiments with resolved immune cell types would be of interest to compare the proteomic differences in macrophages from TEN and MPR patients.

This analysis demonstrates that limpa is completely compatible with existing limma pipelines, allowing any arbitrarily complex experimental design. Leveraging on limma, limpa supports a wide range of analysis tasks ranging from surrogate variable analysis, biological variation modelling, to GO and pathway analysis.

## Discussion

This article introduces limpa for the analysis of label-free proteomics data. The DPC [24] provides a probabilistic model to formally describe the missingness patterns in label-free proteomics data, which can be used to inform the downstream analysis. Based on DPC, we propose a novel statistical approach for protein quantification called DPC-Quant. Precursor intensities are summarised into protein quantities during which missing values are represented by the DPC. An empirical Bayes approach is adopted to borrow information across all the peptides identified and quantified in the experiment. No missing data exist in the protein-level summarisation by DPC-Quant. Accompanying the protein quantification results, DPC-Quant also measures the amount of uncertainty associated with each protein value. Quantification uncertainty is taken into account in DE analysis via a novel limma-style pipeline called vooma. The vooma pipeline generates precision weights by interpolating the global variance trend onto each observation, which are then propagated into the linear models in DE analysis. The variance trend is fitted using a bivariate linear model to capture both the mean-variance trend and the heteroscedasticity induced by uncertainty in protein quantification. Using human mixture data, we observed that both quantification uncertainty and average log-intensity are strong predictors of residual variances (Supplementary Fig. 1).

In our benchmarking analysis, whenever limma was applied, we used the limma-trend approach where protein-wise variances are squeezed towards a global mean-variance trend. We notice that in current proteomics literature, the limma-trend option is often overlooked. It is noteworthy that the limma-trend method is often more powerful than the default for many datasets, especially when a clear mean-variance relationship is present. Our benchmarking is therefore comparing most of the existing methods in a more powerful form than is usually the case.

We demonstrated using a high-quality mixed-species dataset that most of the existing imputation methods gave biased logFC results. In contrast, DPC-Quant provides consistently reliable logFC estimates even for proteins of low intensities with a lot of missing values. Additionally, accuracy in protein quantification depends not only on the precursor-to-protein summarisation, but on the upstream software that quantifies the precursors as well. For the same experiment, if quantified by Spectronaut, we see from the MD plots (Supplementary Fig. 11) that all imputation-based pipelines give systematically biased logFC estimates at low intensities, similar to what was observed on data searched and quantified by DIA-NN. However, although both NA-approaches (NA-MaxLFQ and NA-Top1) performed reasonably well on data processed by DIA-NN (Supplementary Fig. 4), their results were strikingly worse on Spectronaut data (Supplementary Fig. 11). This suggests that the quality of precursor-level quantification, if not more important than the imputation and the protein quantification algorithms, also has the decisive impact on the accuracy of protein quantification, consequently on the reliability of DE results.

The CPXA ECOLI protein in particular had very inaccurate logFC estimates returned by many of the existing pipelines, including v2mnar-MaxLFQ, all the protein-level and NA-based pipelines, and by MaxLFQ-proDA (Supplementary Fig. 11). For all of these pipelines, the protein had extremely high quantified log-intensities in group B, which were driven by one particular precursor for the protein, as shown in Supplementary Fig. 12. Meanwhile DPC-Quant gave more robust results because it weighted all the precursors for that protein more equally (Supplementary Fig. 11).

DPC-Quant has the generality to incorporate prior quality weights into the protein quantification algorithm. In the current implementation, all precursors, in fact all observations, are weighted equally by default in the additive linear model. This means each precursor and each observation contribute equally to the quantification of the protein. One interesting idea for future exploration is to introduce weights to each precursor-level observation through some quality metrics when additive linear model is fitted. Ideally, the quality metric should describe the level of confidence in each precursor identification and quantification, so that those of poor quality can be reliably downweighted in protein summarisation. In this way, it might be possible to retain more peptide quantifications with lower quality metrics in the dataset without compromising the quality of the final quantifications.

limpa’s voomaLmFitWithImputation function is based on the vooma and voomaLmFit functions in limma, all of which can accept any variance predictors as arguments. This means that the vooma pipeline does not always need to be paired with DPC-Quant, and can be used as an independent method. For example, it was shown in Zhu *et al*. (2020) [31] that protein variances can also depend on the peptide-spectrum-match (PSM) count or the number of peptides used in protein quantification for each protein. In fact, such an analysis has been possible directly in limma since version 3.52.0 by inputting the protein-wise PSM or peptide count to the trend argument in eBayes() function. Yet this yields only a univariate trend, and we overlook the mean-variance relationship. With the latest vooma pipelines, one can perform the bivariate variance modelling so that average expression and peptide count of each protein are simultaneously taken into account. This can be done simply by inputting the PSM or peptide count to the predictor argument of voomaLmFit. The DPC-Quant method described in this article can be viewed as a more developed version of this approach, and the peptide counts are already integrated into the quantification uncertainty if DPC-Quant is used.

The use of mixed-species samples is a common way to create calibration datasets for the purpose of evaluating instruments, workflows and software tools in label-free proteomics. Proteins from different species are mixed at fixed proportions, so that the expected logFC of each protein is known based on its species. Compared to the conventional spiked-in design where only a small number of proteins (e.g., using the Universal Proteomics Standard sets) are DE between groups, the mixing species design enables a more comprehensive evaluation on quantification given more changing proteins covering a wider dynamic range. A key application of the mixing species design is for evaluating the coverage and quantification accuracy of proteins by different MS instruments, methods and/or software tools [45, 9, 46]. However, care must be taken when using mixed-species experiments for evaluation of DE methods.

One concern about mixed-species calibration datasets is whether they are able to realistically represent real biological datasets. On the one hand, typically only technical replications are created, so that biological variations are not considered. On the other hand, proteins of different species exhibit disparate variation patterns. In Supplementary Fig. 13A, we visualise the distributions of precursor- and protein-wise standard deviations of each species in the mixed-species dataset processed by DIA-NN. Standard deviation is calculated ignoring the missing values for both precursor and protein levels, where protein quantification is performed using the NA-MaxLFQ pipeline. We observed that *E. coli* and yeast proteins are more variable across samples compared to the background on both precursor and protein levels. This is somewhat expected since they are genuinely DE between groups. However, we are also concerned by the bimodal distributions in the standard deviation of *E. coli* and yeast precursors.

Such observations have great implications on the precision of DE analysis, which relies on variance modelling. Both limma-trend and vooma pipelines assume proteins measured in the same experiment follow a global mean-variance trend in order to borrow information between proteins for better statistical power. Naturally one question arises, that is, whether it is still feasible to assume a global mean-variance trend when proteins from difference species clearly have distinct variance patterns. As demonstrated in Supplementary Fig. 13B, *E. coli* and yeast proteins had distinct mean-variance relationships compared to human proteins. The same was true for both quantification uncertainty-variance relationship (Supplementary Fig. 13C) and the vooma variance trend (Supplementary Fig. 13D). We can certainly perform a bespoke DE analysis for this dataset by modelling a separate variance trend for each species. Yet this defeats the idea of benchmarking where the methods are supposed to be evaluated under a realistic and representative setting of real biological experiments. Generally, the mixed-species design is a useful resource for evaluating the accuracy of precursor and protein quantification. However, when benchmarking DE methods, careful exploratory analysis must be carried out before drawing the conclusions.

We have also demonstrated the limpa pipeline on a public single cell proteomics dataset. The DPC curve had a slope of *β*_1_ = 0.85 (Supplementary Fig. 8B), suggesting a strong intensity-dependent trend in the missing values. In the presented analysis, the global DPC fitted on all cells from different groups was applied in protein summarisation by DPC-Quant. Supplementary Fig. 8C displays the group-wise DPCs, and we observed slightly different values in the estimated DPC parameters between groups. We also observed in the pure tissue lysate dataset that the tissue-specific DPC curves had much steeper slopes compared to the global curve (Supplementary Fig. 7C,D). It is worthwhile to explore whether applying group-specific DPCs in DPC-Quant improves protein quantification and consequently the DE results for experiments with a discrete group design. Such a strategy will be particularly beneficial for studies where distinct samples are being analysed.

The limpa software provides a complete computational pipeline for analysing MS-based label-free proteomics data. limpa works seamlessly with all existing limma pipelines, allowing any arbitrarily complex experimental design and a wide range of downstream tasks such as gene ontology and pathway analysis. Through extensive evaluation and benchmarking comparisons, we show that limpa outperforms all existing pipelines for label-free proteomics data, especially when missing values are a concern.

## Methods

### DPC estimation assuming complete data are normally distributed

#### Model assumptions

Previously in Li and Smyth (2023) [24], we gave the mathematical derivation of detection probability curve (DPC) assuming the observed values in each precursor follow a normal distribution. Alternatively, one can also assume that for each precursor, the complete data, observed and unobserved, are from a normal distribution.

Let *y* be a log-intensity value and *d* be the indicator of detection with *d* = 1 if *y* is observed and *d* = 0 if *y* is missing. We assume the complete expression in each precursor is normally distributed, that is,

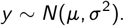

The probability density function of *y* is given by

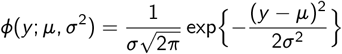

where *µ* and *σ*^2^ are precursor-specific parameters that are assumed to be known.

Based on Li and Smyth (2023) [24], we continute to assume that the detection probability is a logit-linear function of *y*, that is,

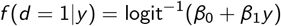

where *β*_0_ and *β*_1_ are the DPC parameters we wish to estimate.

When *y* is missing, i.e., *d* = 0, we have *f* (*d, y*) = *f* (*d* = 0), which is the marginal probability for missing values, and is given by

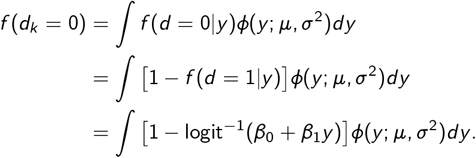

### Estimating the DPC by maximum likelihood

Consider an experiment with *P* precursors and *n* samples. Let *y*_*ji*_ denote the log2-intensity of precursor *j* in sample *i*, where *i* = 1, 2, …, *n* and *j* = 1, 2, …, *P*. The DPC parameters, *β*_0_ and *β*_1_ are estimated by maximising the likelihood

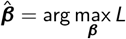

where 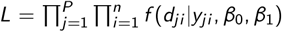. This is equivalent to minimizing minus twice log-likelihood, which can be calculated as

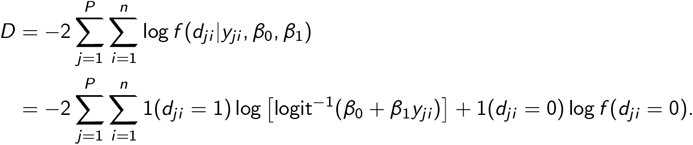

Write *F* () as the cumulative distribution function (cdf) of the standard logistic distribution with location parameter 0 and scale parameter 1. Also write 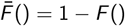 Then

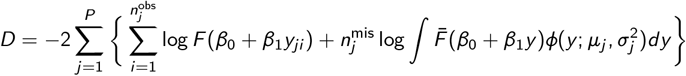

where 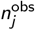 is the number of observed samples and 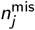 is the number of missing samples in precursor *j*.

The integral in *D* can be can be approximated using Gaussian quadrature. Let *π*_1_, …, *π*_16_ and *z*_1_, …, *z*_16_ be the weights and nodes for 16-point Gauss-Hermite integration, which can be obtained from `gauss.quad.prob(16, dist=“normal”)` using the `statmod` R package. Then we have

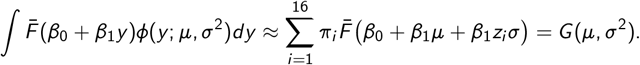

Inserting *G*() for the integral gives

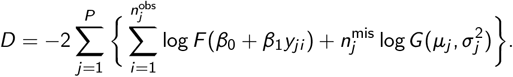

The BFGS algorithm is then used to solve

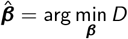

for 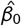 and 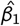. We use the optim() function in R, to which the gradient is supplied. The gradient, 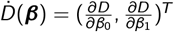, is given by

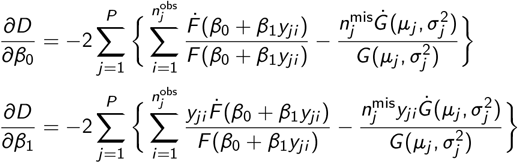

where 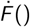 is the first-order derivative of F ().

In the software implementation, estimates for the mean and variance of the normal distribution of each precursor are required, that are *µ*_*j*_ and 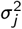 for *j* = 1, 2, …, *P*. We obtain the starting values for these nuisance parameters from imputed precursor-level data output by limpa’s imputeByExpTilt function, which uses a very fast imputation algorithm based on exponential tilting [24]. The DPC coefficients are optimised at given *µ*_*j*_ and 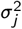 estimates. Given the estimated DPC, we update the nuisance parameters, based on which we update the DPC estimate again. This iterative process is repeated several times until the *µ*_*j*_ and 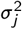 estimates stabilise.

We call this the complete-normal DPC, and is implemented as the dpcCN() function in limpa. The DPC model proposed in Li and Smyth (2023) [24] is referred to as the observed-normal DPC. In real biological datasets, the observed-normal DPC provides a practical approximation of the complete-normal DPC.

### Mathematical derivation of DPC-Quant

#### Model assumptions

Consider a protein with *N* = *p* × *n* observations, *p* being the number of precursors mapped to this protein and *n* being the sample size. For simplicity, use *y*_*k*_ instead of *y*_*ji*_ to denote the log-intensity of sample *i* in precursor *j*, where *i* = 1, 2, …, *n* and *j* = 1, 2, …, *p*. Let *d*_*k*_ be the indicator of detection with *d*_*k*_ = 1 if *y*_*k*_ is observed and *d*_*k*_ = 0 if *y*_*k*_ is missing. Assume

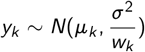

where *σ*^2^ is known and *w*_*k*_ is the observational weight for *y*_*k*_. In practice, *σ*^2^ is a protein-wise nuisance parameter. The probability density for *y*_*k*_ is then given by

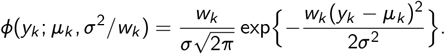

Note that, the normality assumption is made on complete data, where *y*_*k*_ can be a missing value (NA) in the data matrix.

The joint density of detection and log-intensity is denoted as *β* (*d*_*k*_, *y*_*k*_), and when *y*_*k*_ is observed, i.e., *d*_*k*_ = 1, we have

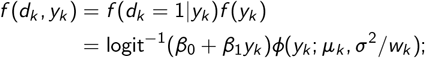

where *f* (*d*_*k*_ = 1|*y*_*k*_) = logit^−1^(*β*_0_ + *β*_1_*y*_*k*_) is the probability of *y*_*k*_ being detected, and is defined by the DPC.

When *y*_*k*_ is missing, i.e., *d*_*k*_ = 0, we have *β* (*d*_*k*_, *y*_*k*_) = *β* (*d*_*k*_ = 0), which is the marginal probability for missing values, and is given by

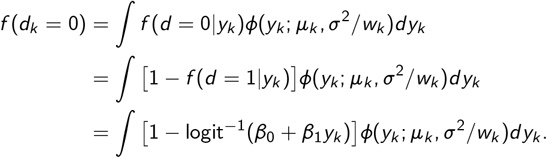

#### Additive linear models for protein quantification

For each protein, we fit the additive model where *µ*_*k*_ = sample_*i*_ + precursor_*j*_, or formally,

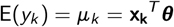

where ***θ*** is a vector of unknown coefficients representing sample and precursor effects; and **x**_*k*_ is a binary vector indicating which sample and precursor *y*_*k*_ is from. Use *γ*_*i*_ to represent the sample effect in sample *i* and *δ*_*j*_ for precursor effect in precursor *j*, we then have

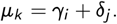

The sum-to-zero contrast coding is used for precursor effects so that *δ*_*j*_ represent the baseline differences among the precursors and 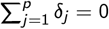. The sample effects, *γ*_*i*_, can then be interpreted as the mean log-intensity in each sample. The last precursor parameter *δ*_*p*_ is redundant so

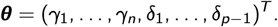

#### Hierachical model

We assume an uninformative prior on the precursor effects, denoted by ***δ***_*g*_ = (*δ*_*g*1_, …, *δ*_*gp*_)^*T*^ for the *g* -th protein. For the sample effect *γ*_*i*_, assume

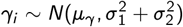

and

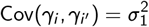

for any *i* ≠ *i* ^′^. Such a prior is assumed as we expect samples to be correlated through being measured on the same protein; and variation in experssion tends to be smaller between samples for the same protein than between proteins for the same sample. The prior for average expression is thereby

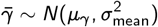

where 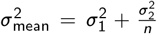, which is the variance in mean expression across all proteins. If **c**_*j*_ is a *n*-vector such that 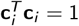 and 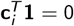, then 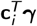 is independent of 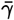 and

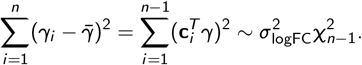

where 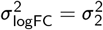. We can then write out the kernel for minus twice the log prior as

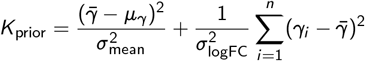

where 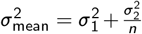 and 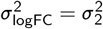. The first term represents between-protein variation and the second within-protein variation. There are three hyperparameters arising from this prior information, which are 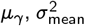 and 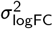, or equivalently, 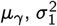 and 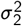.

The posterior distribution for ***θ*** is thus

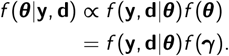

For each protein, we estimate 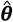 to obtain the protein-level summary, 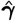, by maximising this posterior.

#### Maximum posterior estimation of model coefficients

Sample and precursor coefficients are estimated by maximising the posterior distribution

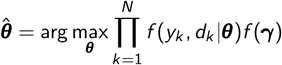

which is equivalent to minimizing negative twice the log-posterior, that is,

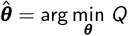

where

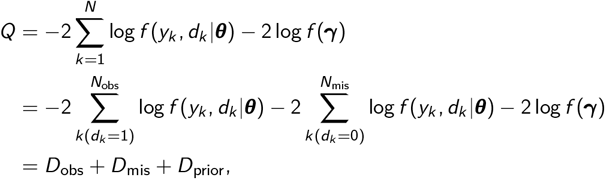

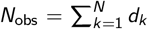 is the number of observed log-intensity values on the precursor level and 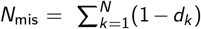 is the number of missing values in this protein. We see that *Q* is partitioned into three independent parts, which are the deviance arising from observed values *D*_obs_, the deviance arising from missing values *D*_mis_ and the deviance of the multivariate prior *D*_prior_. The kernel of the observed deviance is given by

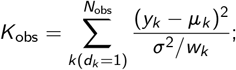

and the deviance corresponding to the missing values is

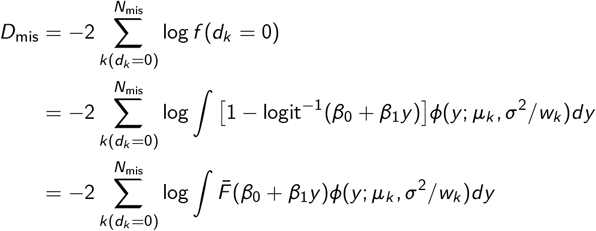

where *ϕ* () is the normal probability density function (pdf) and 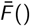 is the right tail cdf of the logistic distribution, i.e., 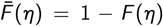 where *F* () is the cdf of the standard logistic distribution with location parameter 0 and scale parameter 1.

As before, we approximate the integral in *D*_mis_ using Gaussian quadrature. Inserting *G*() for the integral gives

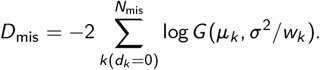

The optimization objective, *Q*, can then be written as

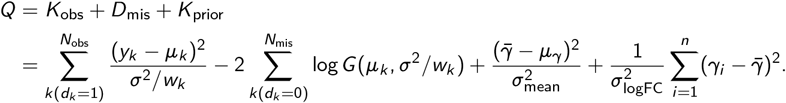

Let 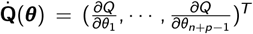 be the score vector. The *s*-th element of 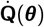 is the first-order derivative of *Q* with respect to *θ*_*s*_, i.e.,

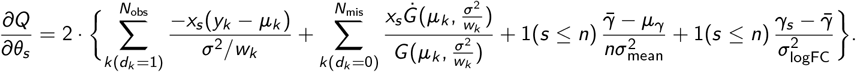

Note that for the *D*_mis_ term, we are directly differentiating the nodes of the Gaussian quadrature approximation.

The maximum posterior estimates for ***θ*** can then be obtained by solving the score equations

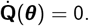

In practice, these score equations must be solved numerically, for which the Hessian matrix is needed. Write 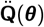 for the Hessian with elements

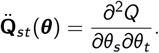

For *s ≠ t* we have

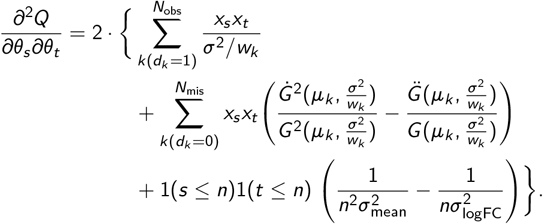

On the diagonal of 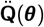, we have *s* = *t*, where

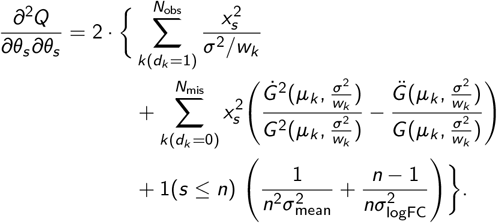

The standard error associated with 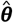 can then be calculated as the square root of the diagonal elements of 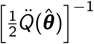 i.e.,

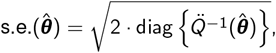

which quantifies the precision of the estimate of the coefficient. We take the standard error of the maximum posterior estimate of each sample effect, s.e.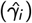, as a measure of uncertainty in the protein-level summary in sample *i* for this protein.

#### Estimation of hyperparameters and nuisance parameters

The maximum posterior estimation for protein quantification depends on three hyperparameters, 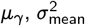 and 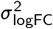. It also depends on 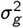, which is the variance of precursor-level log-intensities in the *g* -th protein, and 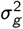 is a protein-wise nuisance parameter for *g* = 1, …, *G*. We use the empirical Bayes approach to estimate the hyperparameters. Given the prior distribution is assumed on complete data, we need to estimate both hyperparameters and the nuisance parameters from complete data with no missing values. As a practical strategy, we obtain approximate complete data by imputing precursor-level missing values with limpa’s imputeByExpTilt function, which uses exponential tilting as described in Li & Smyth (2023) [24].

Specifically, *µ*_*γ*_ is the mean of average protein expression, which describes the global protein expression level in data. We estimate *µ*_*γ*_ by the mean of average log-intensities of precursors, that is,

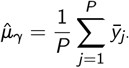

where 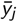 is the average log-expression of precursor *j*. Similarly, 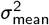 is the variance of average protein expression which depicts the between-protein variation. We get an estimate for 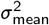 by the variance of average log-intensities of precursors, that is,

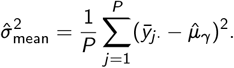

The last hyperparameter 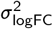 represents the within-protein variation. We get an estimate by taking the *q*-quantile of all within precursor variances, i.e.,

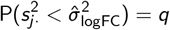

where q ∈ [0, 1] and 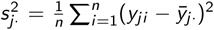 is the sample variance of precursor *j, j* = 1, …, *P*. The default is set to *q* = 0.9 in `limpa`.

To estimate 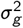, we fit the same “sample + precursor” model for protein quantification on completed data. That is, for each protein, we fit

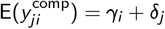

where 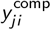 is the log-intensity of precursor *j* in sample *i* in the completed data. Coefficients are estimated by the maximum likelihood approach, and the residual variance from each protein-wise model is extracted and used as an estimate of 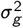 for protein *g*.

The DPC-Quant method is implemented in the dpcQuant function of the limpa package. The function inputs the matrix of peptide-level log-intensities, estimates the necessary hyperparameters, maximises the posterior distribution, and outputs an expression object containing the matrix of protein-level log-intensities.

### Precision weights to account for quantification uncertainty in DE analysis

Variance modeling and weight interpolation are implemented in limpa’s voomaLmFitWithImputation() function, which uses a generalization of the precision weights approach first proposed by Law *et al*. (2014) [38]. A bivariate linear model is used to summarise how the variance in protein intensities depends jointly on average expression and the quantification uncertainty.

We first fit the regression model for DE analysis, i.e.,

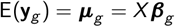

on the protein intensities estimated by DPC-Quant. Fitted values and the residual standard deviation can be obtained where

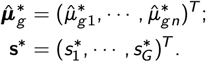

To simplify the notation, we will use *ξ*_*gi*_ for log [s.e.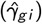] which is the log-uncertainty. We fit the bivariate linear model on protein-wise square root residual standard deviations such that

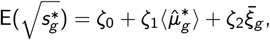

where

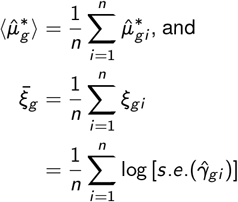

are the average log-expression and the average log-uncertainty in protein *g* respectively. For each protein *g*, we get the fitted square root residual standard deviation as

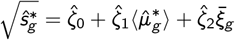

where in general we expect 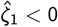 and 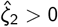, as greater residual variances are typically associated with low abundance proteins of greater quantification uncertainty.

A lowess curve,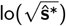 is then computed using these fitted values to form a local smoothing function which describes the variance trend, where 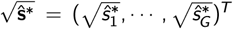. For each protein observation *y*_*gi*_, the square root residual standard deviation is estimated as

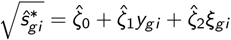

and we use the inverse of interpolated residual variance as its precision weight *v*_*gi*_ such that

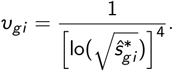

limpa’s voomaLmFitWithImputation() function calculates precision weights and uses the weights to fit linear models while automatically estimating sample weights and block correlations if those are specified. It also adjusts the residual variances and residual degrees of freedom to avoid using variance contributions from groups that are entirely imputed for any protein. The dpcDE() provides an easy user-interface, extracting standard errors from the dpcQuant() output and inputing them into voomLmFitWithImputation(). The fitted model object produced by dpcDE() then enters into standard limma pipelines, for example with eBayes() and topTable().

### Datasets

#### Mixed-species dataset

Hybrid proteome samples were generated by mixing yeast, Escherichia coli (*E. coli*) and human lysates and analysed by SWATH-MS [45]. Yeast and *E. coli* lysates were added to a constant human background at two different ratios. We consider the HYE110 dataset (TripleTOF 6600; 64-variable-window acquisition) where for both mixing conditions, technical triplicates were generated (*n* = 6), and a summary of the mixing proportions is given in Fig. 2A. The expected log 2 fold change comparing Group B versus Group A, log_2_(B*/*A), is log_2_(67*/*67) = 0 for human proteins, log_2_(30*/*3) = 3.3 for *E. coli* proteins and log_2_(3*/*30) = −3.3 for yeast proteins. The original data were published accompanying the LFQbench software [45], from which details on sample preparation and data acquisition methods are available. The study is accessible from the PRIDE partner repository [47] with the dataset identifier PXD002952.

We downloaded the precursor-level data searched by DIA-NN v1.6.0 on the original MS raw files for the HYE110 dataset under the library-based mode published in [9]. Precursor-level output is obtained from the DIA-NN report, and the log 2-transformation is applied to precursor intensities. The total number of detected precursors is 42, 147 in the report. During pre-processing, we first filter out observations with a ‘Q.Value’ or ‘Protein.Q.Value’ *>* 0.01. We also filter out non-proteotypic precursors and protein groups identified by a single precursor (i.e., *single precursor protein groups*, or *single precursor proteins* in this thesis). In the DIA-NN report, when a precursor is shared by more than one protein, the proteins are concatenated into a new protein group and output as “Protein A;Protein B;Protein C”, etc. We call such a protein group the *compound protein group*, and filter them out in pre-processing. After filtering, the number of unique precursor IDs is 38,081, mapped to 4,272 protein groups. The numbers of precursors and proteins detected for the three species are summarised in Supplementary Fig. 3D. There are approximately 27.6% missing values in the precursor-by-sample data matrix after pre-processing.

We also downloaded the output for the same experiement searched by Spectronaut 11 from [9]. Each elution group is used as a precursor ID. The total number of detected precursors is 46,045 before filtering. Observations with a ‘EG.Qvalue’ *>* 0.01 are filtered out. Raw precursor intensities that are equal to 1 are set back to `NA`. Precursors mapped to single precursor proteins and compound protein groups are also filtered out. The log 2-transformation is then applied to precursor intensities. After filtering, the number of unique precursor IDs is 36,862 which are mapped to 4,187 proteins. The numbers of precursors and proteins detected for the three species are summarised in Supplementary Fig. 10D. The overall missingness in precursor-level data is 28.7% after pre-processing, similar to the DIA-NN search but with slightly fewer precursor and protein IDs. The observed-normal DPC was used in DPC-Quant.

#### Human mixtures with paired pure tissue lysates

Total protein lysates from normal human adult bladder, frontal lobe and cerebellum total protein (BioChain) were diluted to 1 mg/mL and mixed at proportions summarised in Fig. 3A. Four technical replicates were generated for each mixing group, yielding a total of 24 samples. In addition, we also processed bladder, frontal lobe and cerebellum unmixed individually. Five technical replicates were generated from each pure tissue lysate with a total sample size of 15.

Protein was digested with trypsin and lys-C via S-trap (Protifi). Briefly, 20 *µ*g of protein per sample were reduced and alkylated with 10 mM TCEP (Merck) and 40 mM chloroacetamide (Merck) and incubated at 55^°^ C for 20 minutes. Following acidification with phosphoric acid, precipitation and trapping onto micro S-trap columns, samples were digested with 1 *µ*g each of SOL-u trypsin (Merck) and Lys-C (Wako) at 47°C for 90 minutes. Peptide clean-up was achieved with SDB stage tips (GL Bioscience) and were loaded on a 15 cm IonOpticks column which was maintained at 50°C using a column oven. A Neo Vanquish (Thermo) was directly coupled online with the mass spectrometer (Astral Thermo) and peptides were separated with a binary buffer system of buffer A (0.1% formic acid (FA)) and buffer B (80% acetonitrile plus 0.1% FA), at a flow rate of 400 nL/min. The gradient started at 2% B and increased to 34% in 30 minutes before increasing to 100% within 0.1 minutes and held for 3 minutes prior to returning to 2% and re-equilibrated. The mass spectrometer was operated in positive polarity mode with a capillary temperature of 275°C.

DIA acquisition consisted of an MS1 scan (*m/z* = 380–980) with an AGC target of 5 × 10^6^ and a maximum injection time of 5 ms (*R* = 240,000). DIA scans were acquired with the Astral detector with an AGC target of 8 ×10^4^ and a 3 ms maximum time. Fragmentation occurred in the HCD cell with a normalised stepped collision energy of 25% and the spectra were recorded in profile mode. 199 non-uniform DIA windows across *m/z* = 380–980 were collected with a maximum injection time of 3 ms and a 0.6 s loop control which achieved an average of 5 data points per peak.

Raw MS data were searched by DIA-NN (1.8.1) in library-free mode with match-between-run (MBR) enabled for both experiments. Precursor-level data from the second-pass search are used in our analysis. Observations with the ‘Global.Q.Value’ or the ‘Lib.Q.Value’ greater than 0.01 are replaced by `NA` as per recommendations by the DIA-NN manual. Non-proteotypic precursors and compound proteins are filtered out. Precursor-level intensities are log2-transformed.

In the human mixture dataset, 108,197 unique precursor IDs are retained for 8,230 protein groups after filtering. The overall proportion of missing values is approximately 15.8%. Propportion of missing values in each sample is shown in Supplementary Fig. 6B. In the pure tissue lysates, 135,528 unique precursor IDs are retained for 9,278 protein groups after filtering. The overall proportion of missing values is approximately 33.5%. Supplementary Fig. 7A visualises the proportion of missing values in each sample. The observed-normal DPC was used in DPC-Quant for both datasets.

Using the pure tissue lysate dataset, we can define tissue-specific proteins by assuming such proteins should be mostly observed in replicates of the corresponding tissue type and mostly missing in the other samples. Specifically, for each protein, we calculate the proportion of precursors that are completely observed in each tissue type. For cerebellum-specific proteins, we require the protein to be completely observed in at least 80% of the precursors in cerebellum replicates, completely observed in no more than 20% precursors in the frontal lobe replicates and not completely observed in any precursor in the bladder samples. Such a criterion led to 71 cerebellum-specific proteins. For frontal lobe-specific proteins, we require the protein to be completely observed in at least 80% of the precursors in frontal lobe replicates, completely observed in no more than 20% precursors in the cerebellum replicates and not completely observed in any precursor in the bladder samples. Such a criterion led to 120 frontal lobe-specific proteins. Cerebellum- and frontal lobe-specific proteins can be used as the true discoveries in DE comparisons between any two groups of the mixture design. For bladder-specific proteins, we require the protein to be completely observed in at least 80% of the precursors in bladder replicates, completely observed in no more than 20% precursors in the cerebellum or frontal lobe replicates. Such a criterion led to 107 bladder-specific proteins. Bladder-specific proteins can be used as the false discoveries to evaluate DE results on the mixture design.

#### Cell cycle proteomes by single cell proteomics

Single-cell proteomes were profiled by the true single-cell-derived proteomics (T-SCP) pipeline as described in [11]. Four cell populations enriched in different cell cycle stages were produced from HeLa cells by drug treatment. Precursor ions in prepared samples were fragmented in the parallel accumulation-serial fragmentation with data-independent acquisition (diaPASEF) mode [48]. The MS raw files were analysed by DIA-NN in the library-based mode [9]. Details on sample preparation, LC-MS/MS analysis and data processing are provided in [11]. Processed data were downloaded from the ProteomeXchange Consortium via the PRIDE partner repository [47] with the dataset identifier PXD024043. Precursor-level output was obtained from the DIA-NN report and cells from the cell cycle experiment were extracted (n = 231). The number of detected precursor species is 10,754. About 60.4% of the data are missing values. We also set intensity values of zero to be missing, which affected only a small number of values and had little impact on the percentage of missing values. A log 2-transformation is applied to the precursor-level intensities.

Precursor-level intensities were filtered by the q-values on the observation level as recommended by the DIA-NN manual. Precursor intensities with either ‘Global.Q.Value’ or ‘Lib.Q.Value’ greater than 0.01 are filtered out. We also filtered out non-proteotypic precursors and precursors that are mapped to the single precursor proteins or the compound protein groups. This leads to a reduction in the number of precursor IDs to 7,904 corresponding to 1,471 proteins. After pre-processing, the overall missing value proportion is 51.2% on the precursor level. The observed-normal DPC was used in DPC-Quant.

#### Immune cell profiling in cutaneous drug reactions by spatial proteomics

FFPE skin biopsies were acquired from patients with maculopapular rash (MPR), drug reaction with eosinophilia and systemic symptoms (DRESS), toxic epidermal necrolysis (TEN) and healthy individuals [15]. CD45^+^ immune cells were profiled by the deep visual proteomics (DVP) workflow [41]. The timsTOF SCP mass spectrometer was operated under the dia-PASEF mode, and MS raw files were analysed by DIA-NN (1.8.0) in library-free mode. Further details on the experiment are available in in Nordmann *et al*. (2024) [15]. Full DIA-NN output was acquired from the authors. Precursor intensities with either ‘Global.Q.Value’ or ‘Lib.Q.Value’ greater than 0.01 are filtered out. Non-proteotypic precursors and precursors mapped to the compound protein groups are also filtered out. After pre-processing, 5,169 unique precursor IDs are retained for 1,316 protein groups. About 40.4% of the data are missing. A log 2-transformation is applied to the precursor-level intensities. The complete-normal DPC was used in DPC-Quant.

## Competing pipelines

In this article, DE analysis is performed on the protein level, so all DE pipelines to be compared consist of two main steps: protein quantification and DE testing. Supplementary Fig. 2 provides a summary of all pipelines included in the evaluation.

For DPC-Quant, the vooma pipeline implemented in dpcDE() is used for DE testing. DPC-Quant combined with vooma is referred to as the limpa pipeline.

For protein quantification, we compare the new method DPC-Quant to MaxLFQ [49], which has been the dominating method for this task. Many proteomics software including Spectronaut and DIA-NN [9] have their own in-built versions of MaxLFQ. However, we have observed non-ingorable disagreement in MaxLFQ intensities output by different implementations searched on the same dataset. Unfortunately, the reason for such discrepency is not trackable as most of these software are not open-source. For consistency, we compute MaxLFQ intensities using the iq R package v1.10.1 [50].

The next question is whether to perform imputation, and if we do, at what level of data using which imputation method (Supplementary Fig. 2A). We divide existing approaches into three categories: methods that perform precursor-level imputation, methods that perform protein-level imputation, and methods that ignore the missing values, i.e., the NA-approaches.

In terms of the imputation methods, k-nearest neighbour (knn) imputation [51, 52]; random forest (RF) imputation [53] and Bayesian principal component analysis (bpca) imputation [54] are considered in the benchmarking, representing MAR-based methods. We also include MNAR-based methods such as MinDet for left-censored missing data where missing values are replaced by the minimum observed value. In our analysis, these methods are applied on both the precursor and the protein levels. Note that knn, RF and bpca are originally designed for microarray data, but have been generically applied to MS-based data [55, 56, 18, 57, 58]. We also examine MNAR-based methods specialized for proteomics data, including msImpute-v2mnar (v2mnar for short) [59] which imputes precursor intensities, and Perseus [60] which imputes protein quantities. With precursor-level imputation pipelines, we impute the missing values on the precursor level, then MaxLFQ summarisation is performed on completed data. With protein-level imputation pipelines, MaxLFQ summarisation is carried out leaving missing values as is, and imputation is performed on MaxLFQ values.

We use the impute matrix() function from the MsCoreUtils R package (1.18.0) on Bioconductor for knn, RF, bpca and MinDet imputation. We implement our own Perseus imputation in R according to [60], where missing values are replaced by drawing random values from a normal distribution (shift = 1.8 and scale = 0.3). The random seed is set to 1234 for all datasets in Perseus imputation. The v2mnar algorithm is implemented in the msImpute R package (1.16.0). It requires all precursors to be observed in at least 4 samples. Depending on the experiment, this filtering step can lead to a significantly reduced number of quantifiable proteins.

We also consider the NA-MaxLFQ and NA-Top1 approaches, where missing values are left as NA throughout the entire pipelines. Top1 is a method for protein quantification where each protein is simply represented by the precursor with the largest average intensity across samples. Top1 is only included as the NA-approach in our analysis as it discards a substantial amount of precusor-level data, which makes imputation an even more challenging task.

DE tests are then performed on the protein-level data. For all existing pipelines, we use the limmatrend approach where protein-wise variances are squeezed towards a global mean-variance trend rather than assuming a constant prior variance [38]. DE analysis is performed using limma v3.63.3, while limma-trend is implemented by setting trend = TRUE in the eBayes() function. The limma-trend option is always adopted whenever limma is used throughout this article.

MSstats v4.14.1 is also benchmarked. MSstats has functionalities to perform both protein quantification and DE testing. Additionally, DE testing can also be performed using limma on the MSstats protein quantification. Such a pipeline is specified as MSstats-limma in the article.

The proDA method does not perform imputation but incorporates the missing value mechanism into their DE testing procedure. It requires protein-level data therefore we test its performance on NA-MaxLFQ data. The pipeline is labeled as MaxLFQ-proDA in this article.

All analyses are performed in R (4.4.3). All software are run under their default settings unless otherwise specified.

## Data Availability

The human mixture dataset is available on request. All other example datasets used in this article are publicly available from the sources cited in Methods.

## Code Availability

The limpa package is freely available from https://www.bioconductor.org/packages/limpa. The source code can be browsed at https://code.bioconductor.org/browse/limpa or can be git cloned from /packages/limpa.

## Funding

This work was supported by the Chan Zuckerberg Initiative (EOSS grant 2021-237445), by the Australian National Health and Medical Research Council (Fellowship 1058892 to GS, Investigator Grant 2025645 to GS, IRIISS to WEHI), by the University of Melbourne (Melbourne Research Scholarship to ML), by CSL Limited (Translational Data Science Scholarship to ML), and by the Victorian State Government (Operational Infrastructure Support to WEHI).

## Acknowledgements

We thank the WEHI Proteomics Facility. We thank Anna Quaglieri, Toby Dite, Mark Condina, Laura Dagley and Andrew Webb for their involvement and support of the pilot experiment for the mixture design. We thank Samantha Emery-Corbin, Olga Vitek and her group, Thomas Cox, Thierry Nordmann, Maria Tanzer and Jerzy Dziekan for the helpful discussions and/or sharing various datasets during the development of limpa package. We particularly thank Jerzy Dziekan for detailed feedback on a draft manuscript.

## Author contributions

ML developed mathematical methods and software, carried out analyses, wrote the paper. SC undertook MS experiments. GS conceived and supervised the project, co-developed mathematical methods and software, co-wrote and finalised the paper. All authors contributed to and checked the final manuscript.

## Competing interests

None.

## Notes

### Competing Interest Statement

The authors have declared no competing interest.

