## Supplementary Figures 1-13 for "Quantification and differential analysis of mass spectrometry proteomics data with probabilistic recovery of information from missing values"

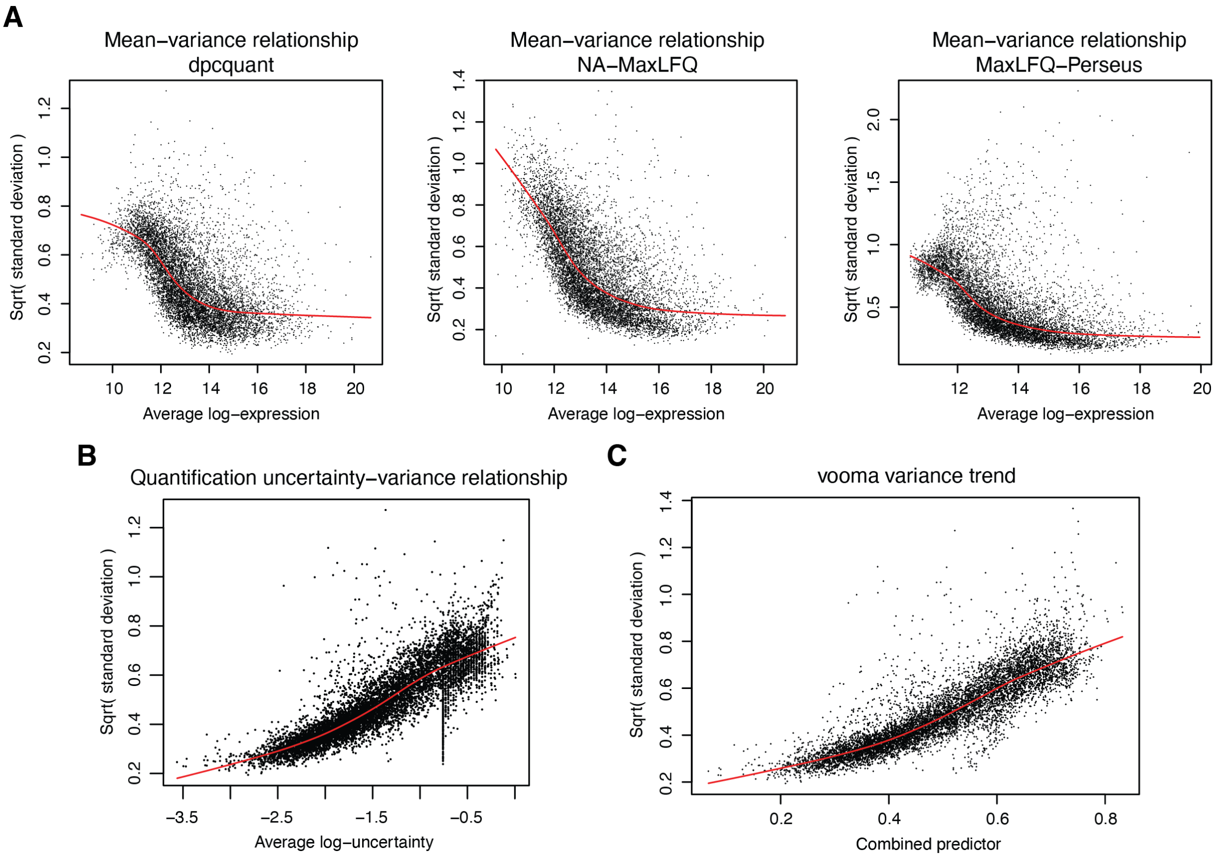


**Figure S1:** On human mixture data, (A) mean-variance relationship, (B) quantification-uncertainty relationship, (C) vooma variance trend on protein quantification by DPC-Quant.

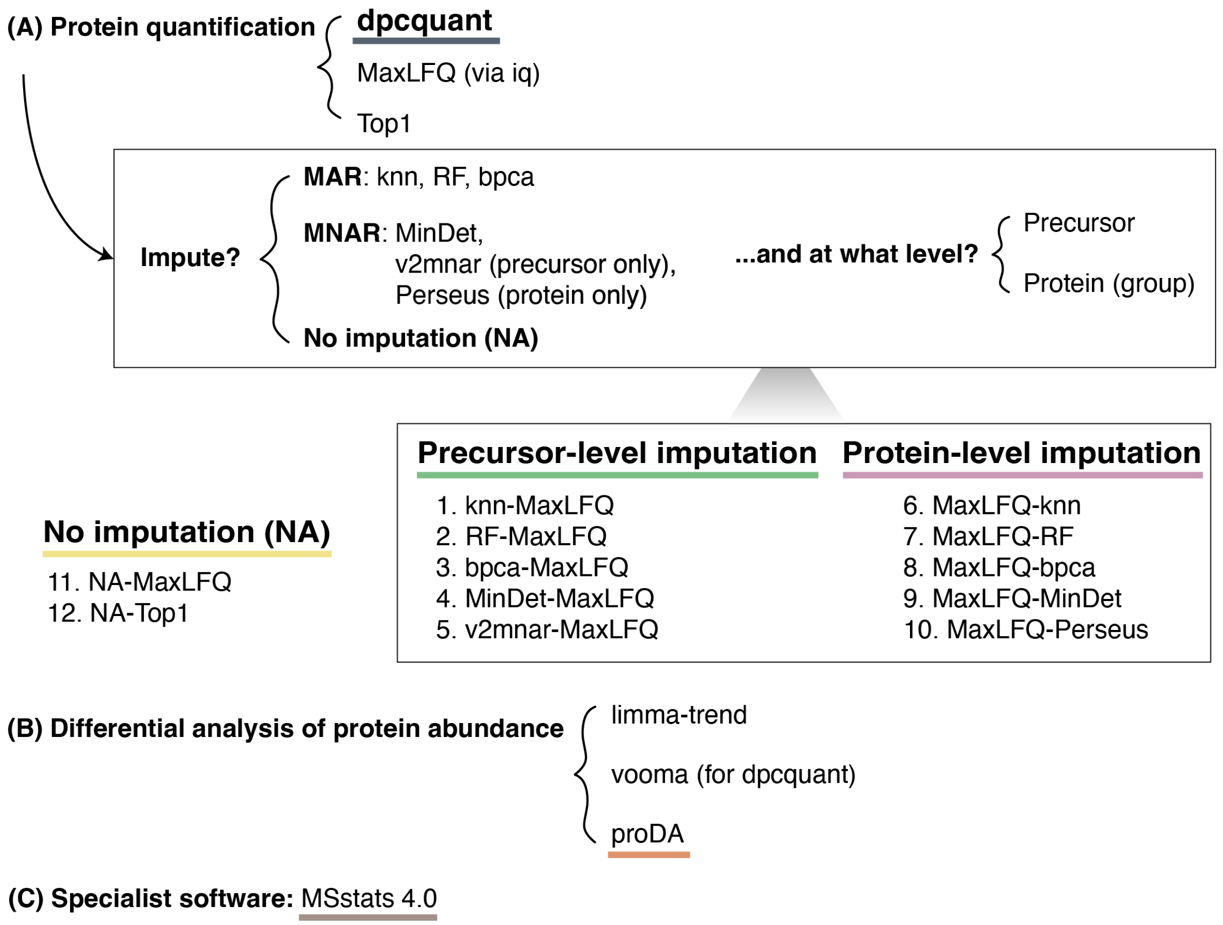


**Figure S2:** Methods and pipelines considered in the benchmarking analyses.

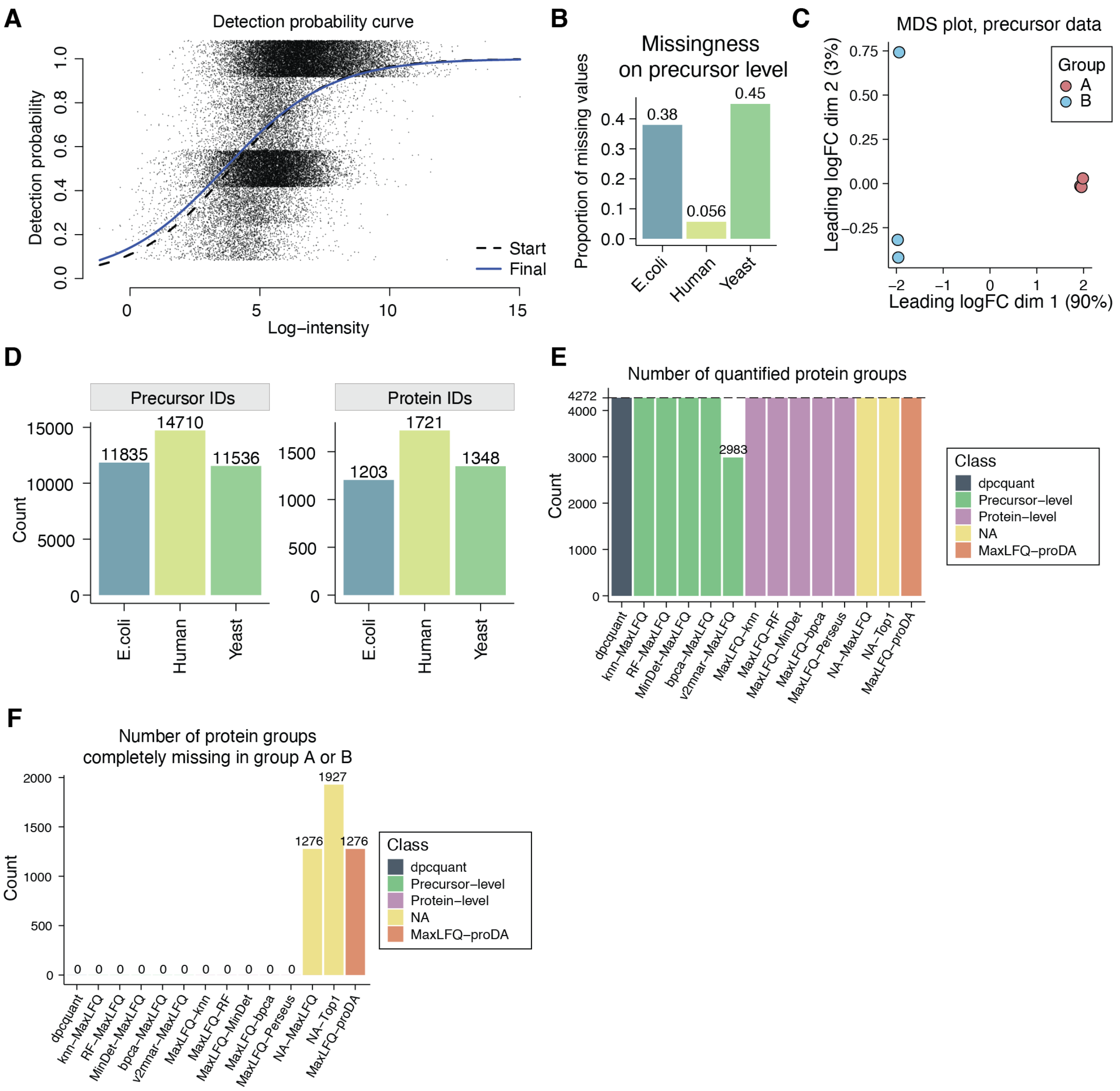


**Figure S3:** Exploration analysis on mixed-species dataset. (A) DPC plot. (B) Proportion of missing values within each species on the precursor level data. (C) MDS plot on the precursor level data. (D) Number of precursor and protein group IDs in each species included in protein quantification. (E) Number of protein groups that were quantified by each pipeline. (F) Number of protein group IDs that are completely missing in all triplicates in group A or B.

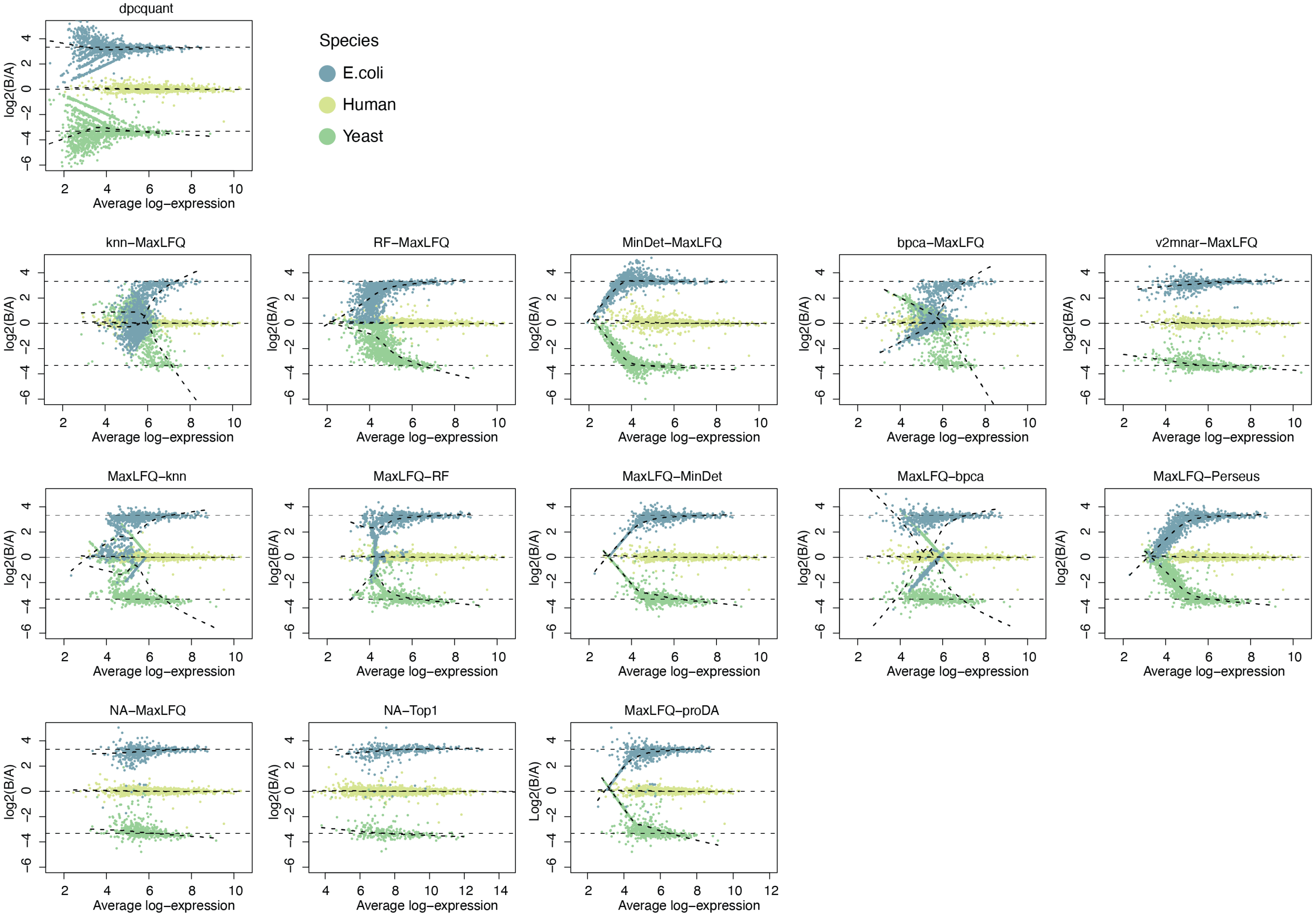


**Figure S4:** Mean-difference (MD) plots comparing group B versus group A on mixed-species data searched by DIA-NN.

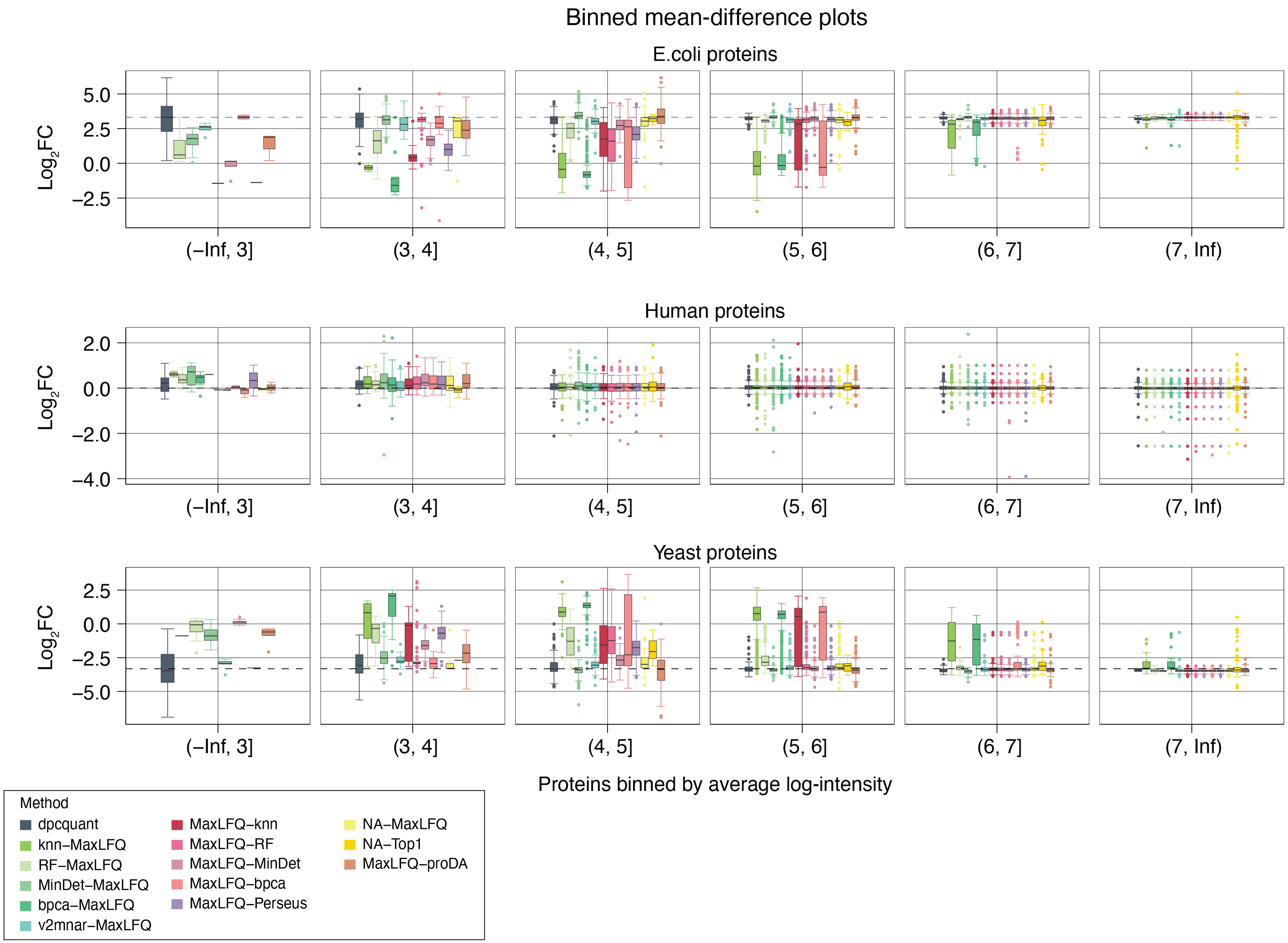


**Figure S5:** Binned mean-difference (MD) plots comparing group B versus group A on mixed-species data searched by DIA-NN.

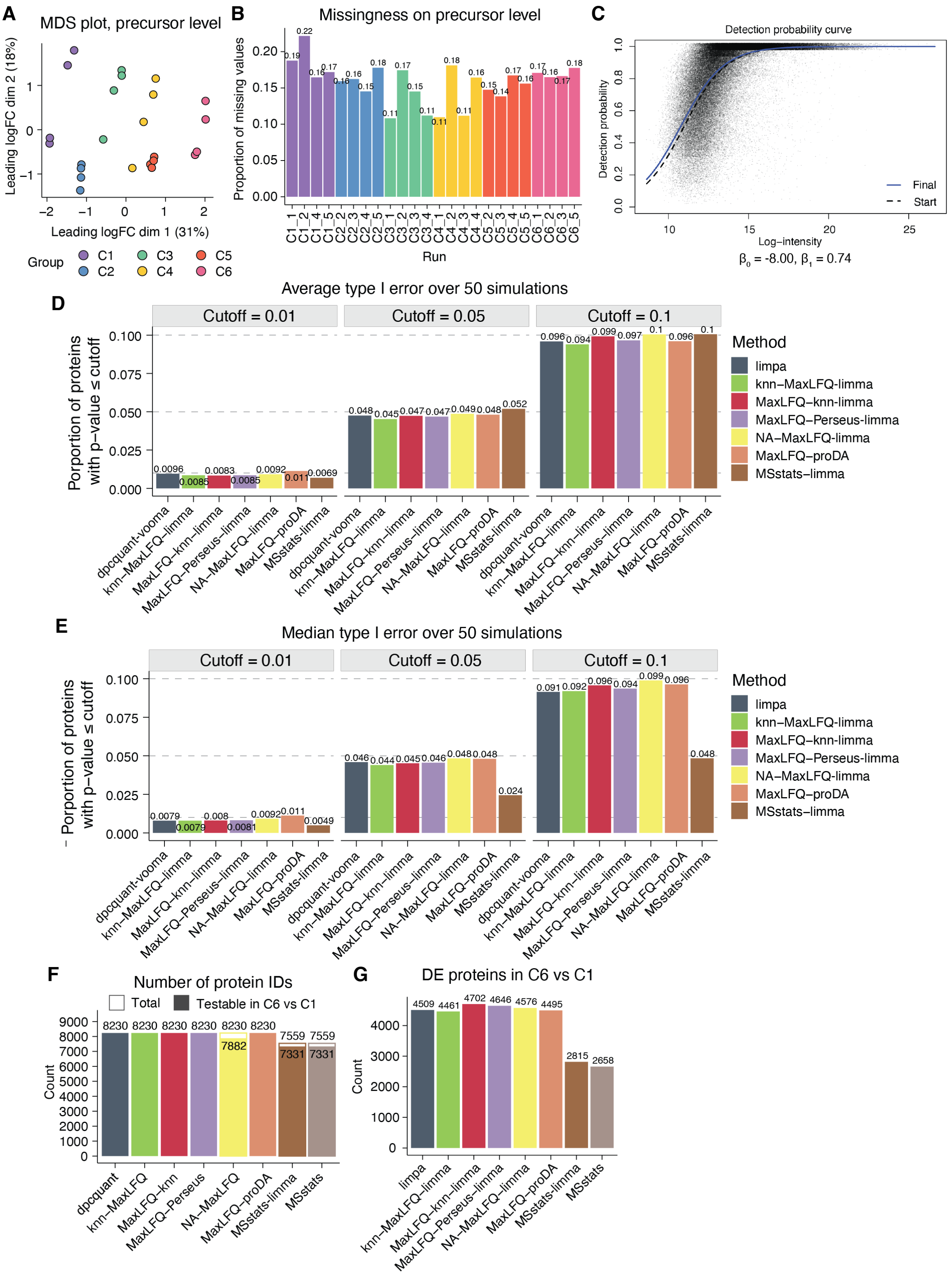


**Figure S6:** Exploratory analysis of the human mixture dataset. (A) MDS plot on the precursor-level data. (B) Proprotion of missing values in each sample on the precursor level. (C) DPC plot. (D) Average type I error rate over 50 simulations of null comparisons at different p-value cutoffs. (E) Median type I error rate over 50 simulations of null comparisons at different p-value cutoffs. (F) Number of protein IDs quantified by each pipeline and the number of proteins with a testable logFC when comparing groups C6 and C1. (G) Number of DE proteins between C6 and C1 at the FDR cutoff of $0.05$ by each pipeline.

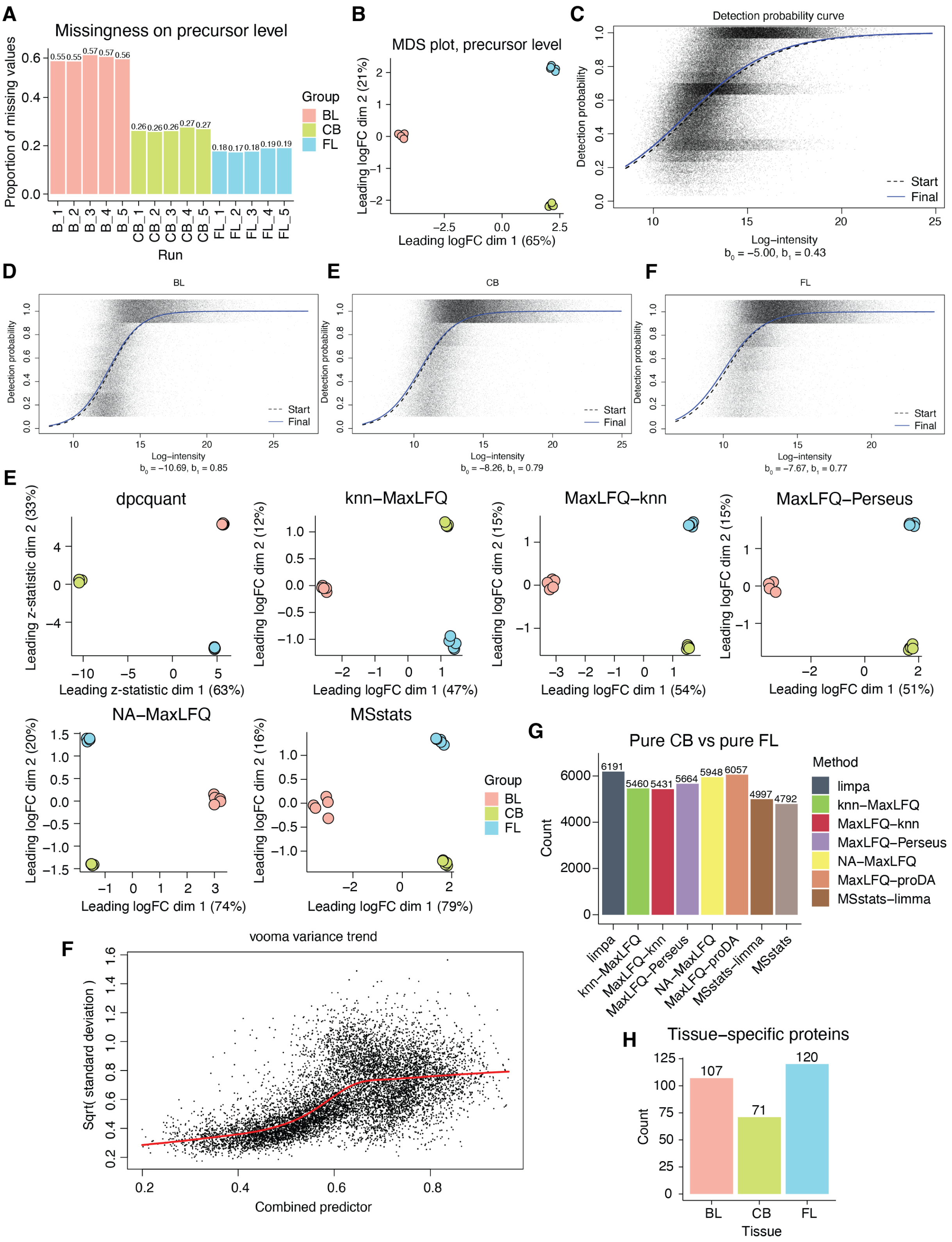


**Figure S7:** Exploratory analysis of the pure tissue dataset. (A) Proprotion of missing values in each sample on the precursor level. (B) MDS plot on the precursor level. (C) DPC plot. (D) DPC estimated within the $5$ replicates of each pure tissue. (E) MDS plot on protein quantification by each pipeline. (F) Vooma variance trend on protein quantification by DPC-Quant. (G) Number of DE proteins comparing the pure cerebellum (CB) samples to the pure frontal lobe (FL) samples by each pipeline. (H) Number of tissue-specific proteins defined by the missing value patterns in each species.

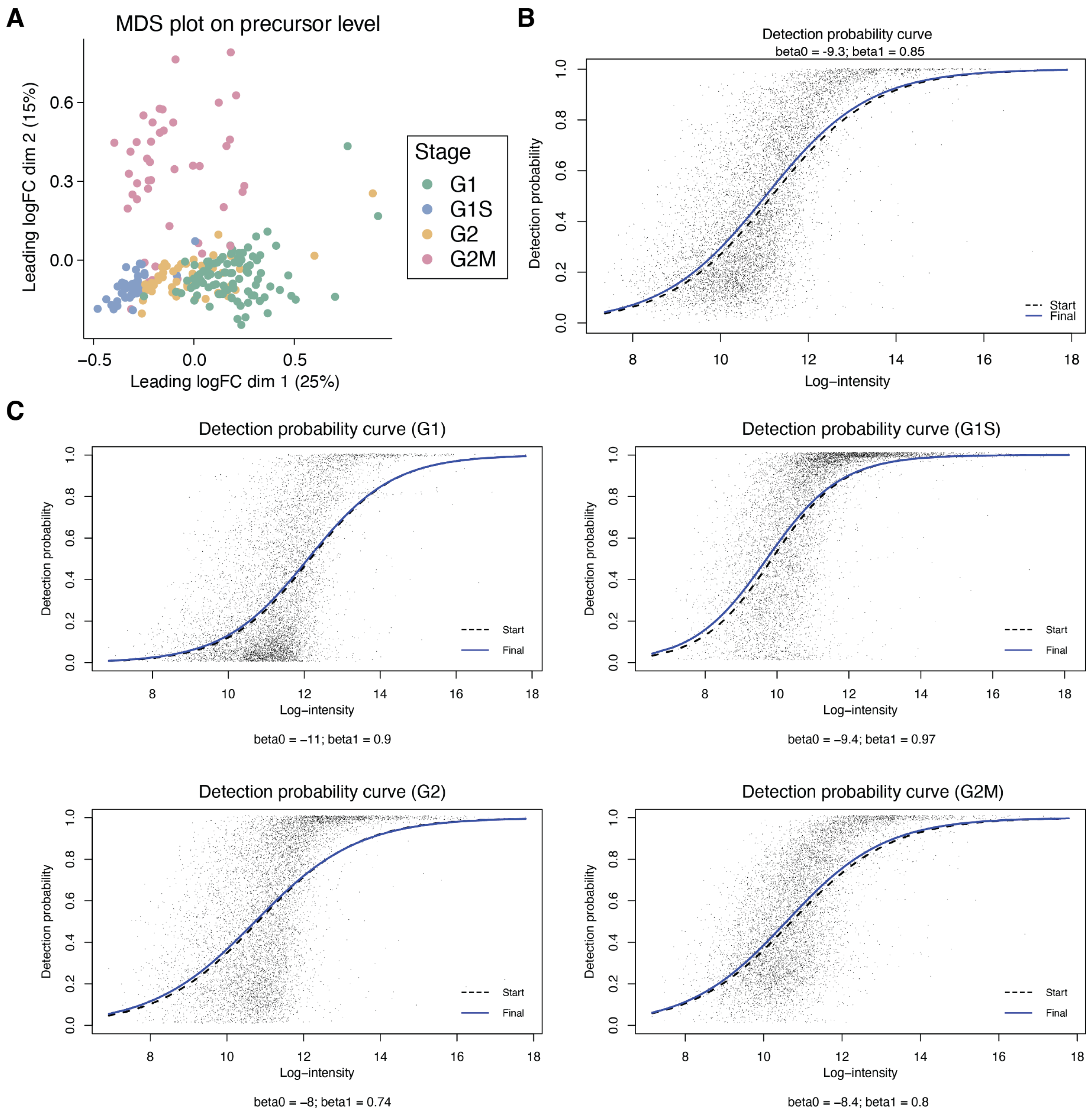


**Figure S8:** Exploratory analysis of the cell cycle dataset. (A) MDS plot on the precursor level. (B) DPC plot on all cells. (C) DPC plots on G1, G1S, G2 and G2M cells, from left to right, top to bottom.

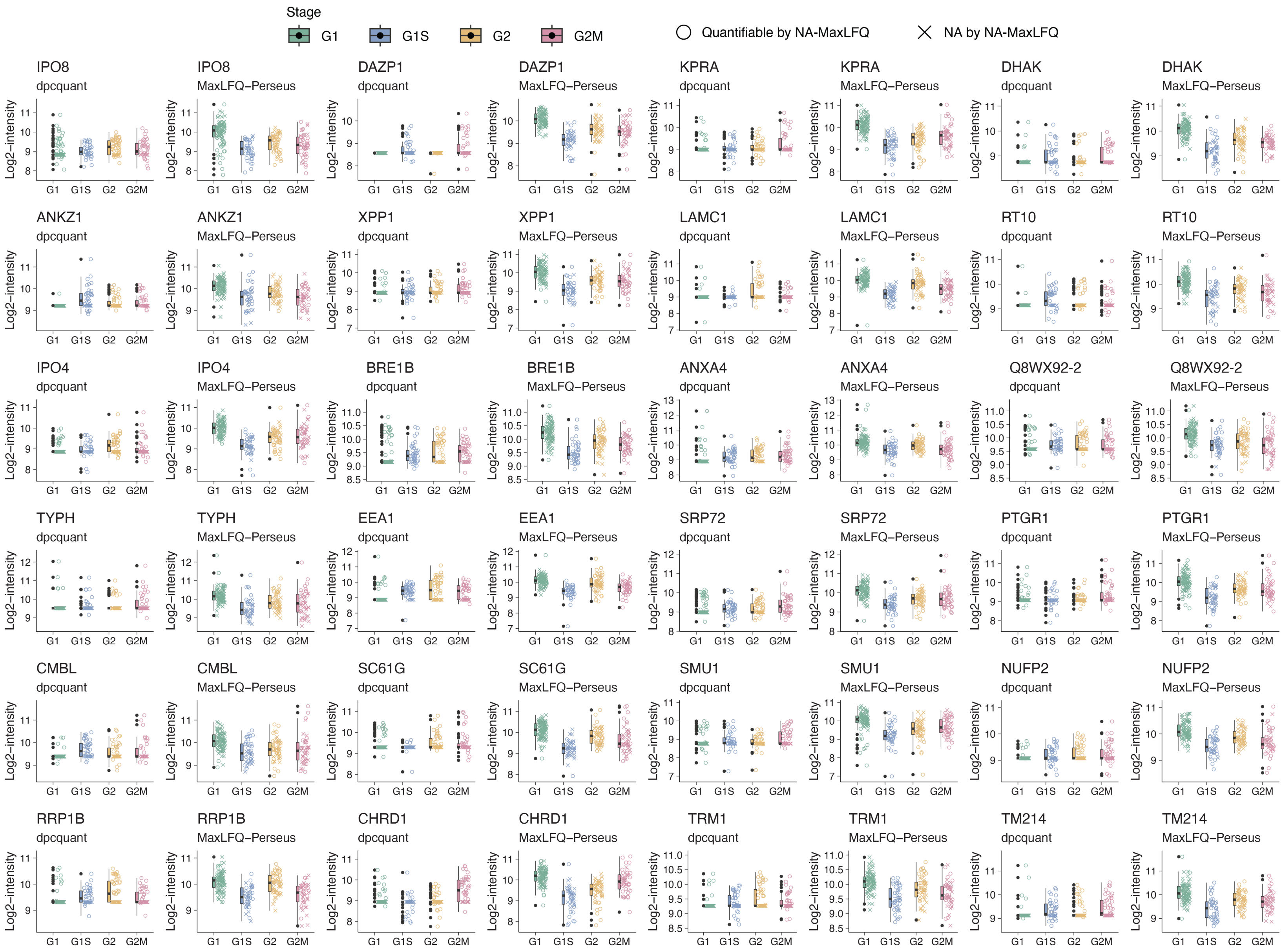


**Figure S9:** Protein-level summary of each cell for protein examples with opposite logFC estimates by DPC-Quant and MaxLFQ-Perseus.

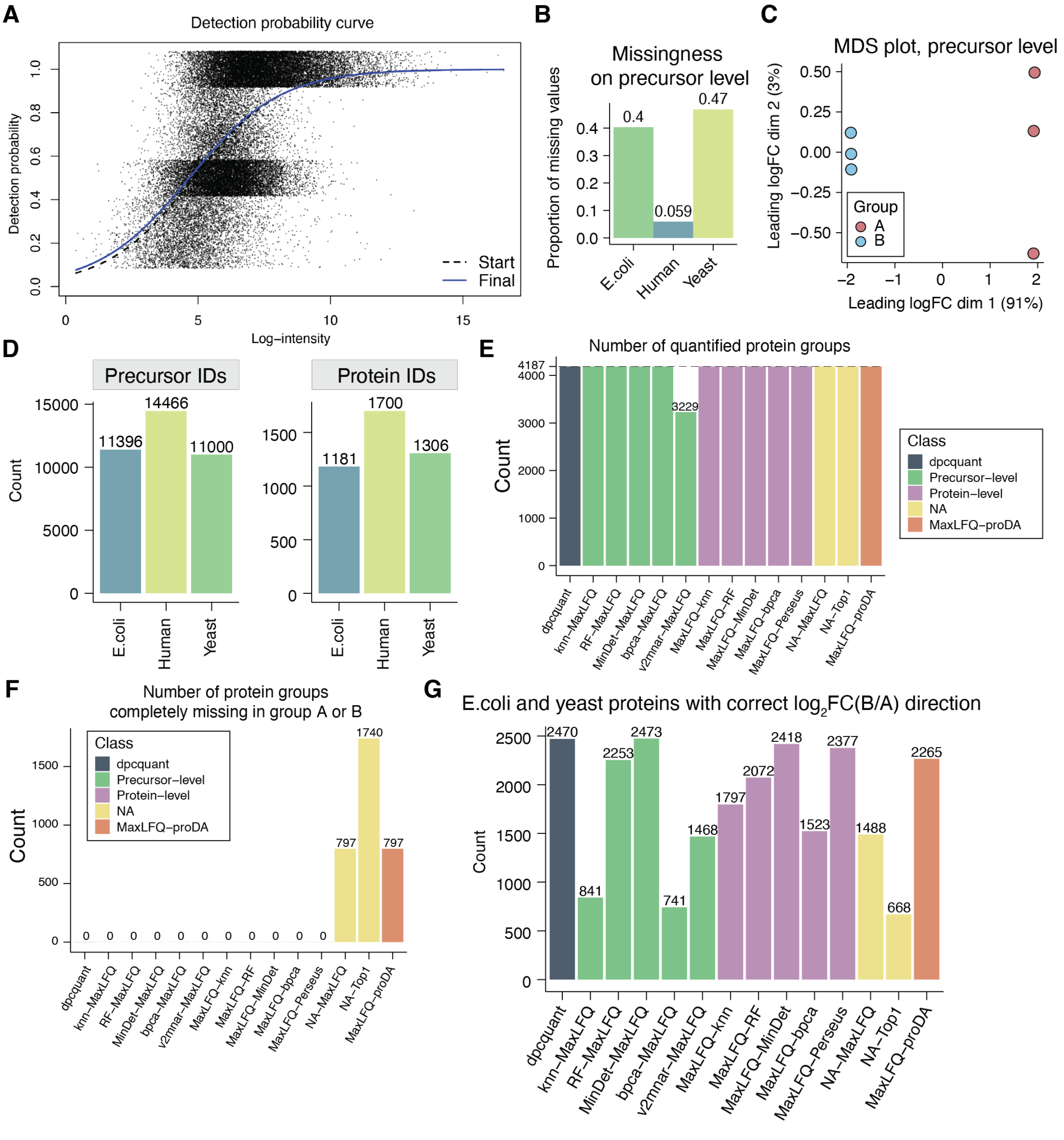


**Figure S10:** Exploratory analysis on mixed-species dataset quantified by Spectronaut. (A) DPC plot. (B) Proportion of missing values on the precursor level of each species. (C) MDS plot on the precursor level. (D) Number of precursor and protein IDs of each species. (E) Number of protein groups that are quantified by each method. (F) Number of protein groups that are completely missing in either group by each method. (G) Number of true DE (yeast and *E. coli*) proteins that have the correct logFC direction.

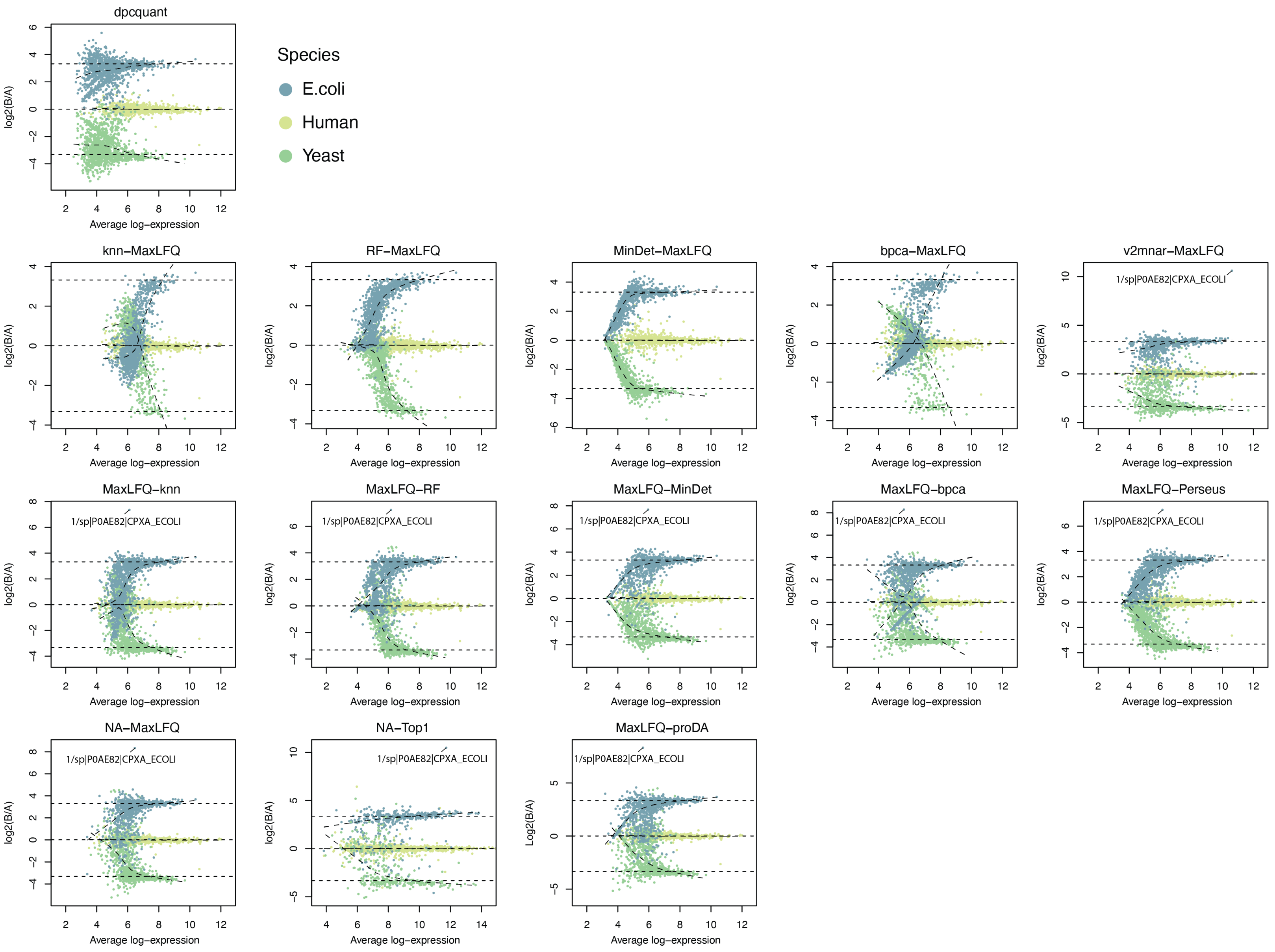


**Figure S11:** Mean-difference (MD) plots comparing group B versus group A on mixed-species data searched by Spectronaut.

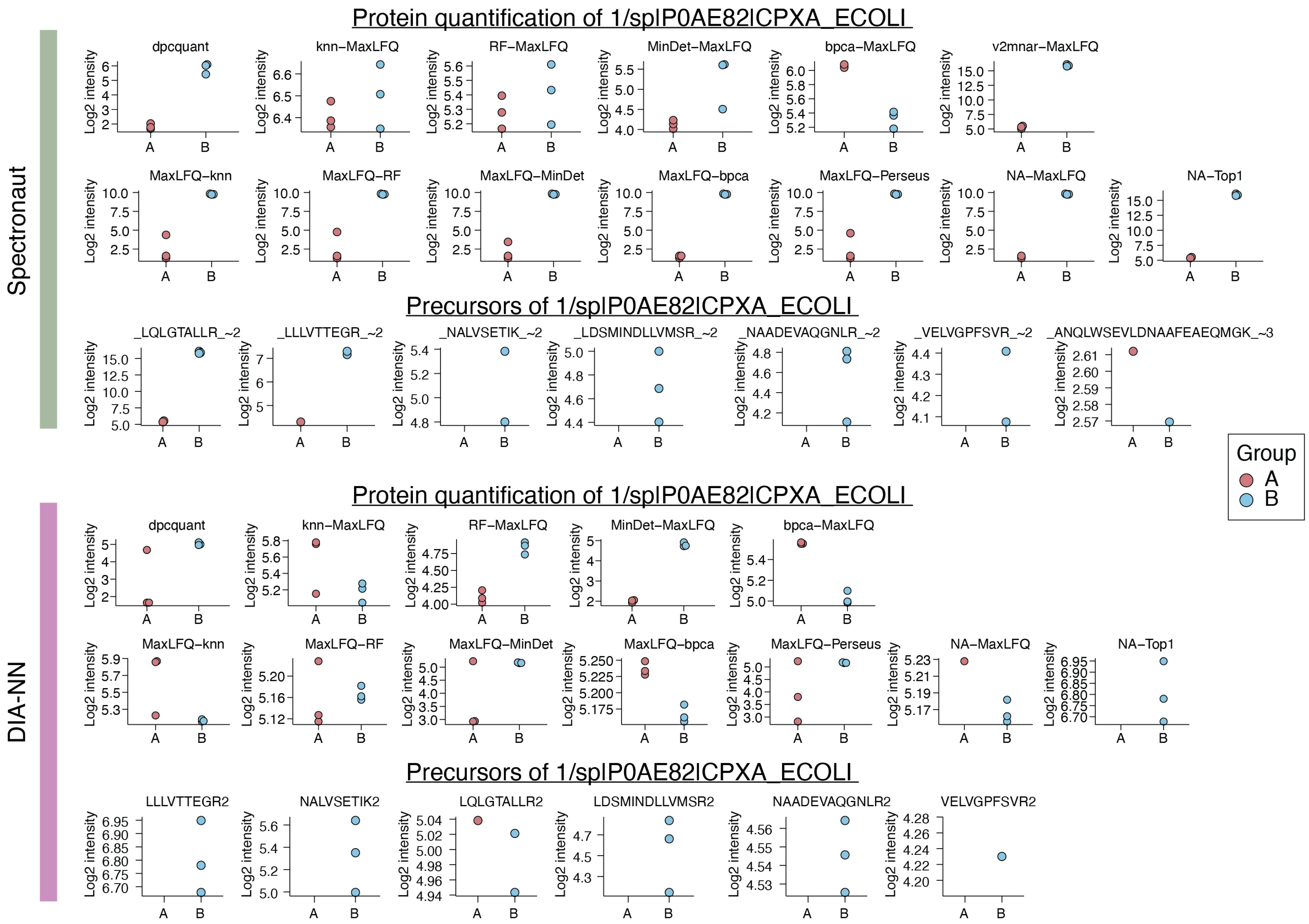


**Figure S12:** Protein-level summary by each pipeline and precursor-level log-intensities of the CPXA_ECOLI protein on mixed-species data quantified by Spectronaut (top) and DIA-NN (bottom).

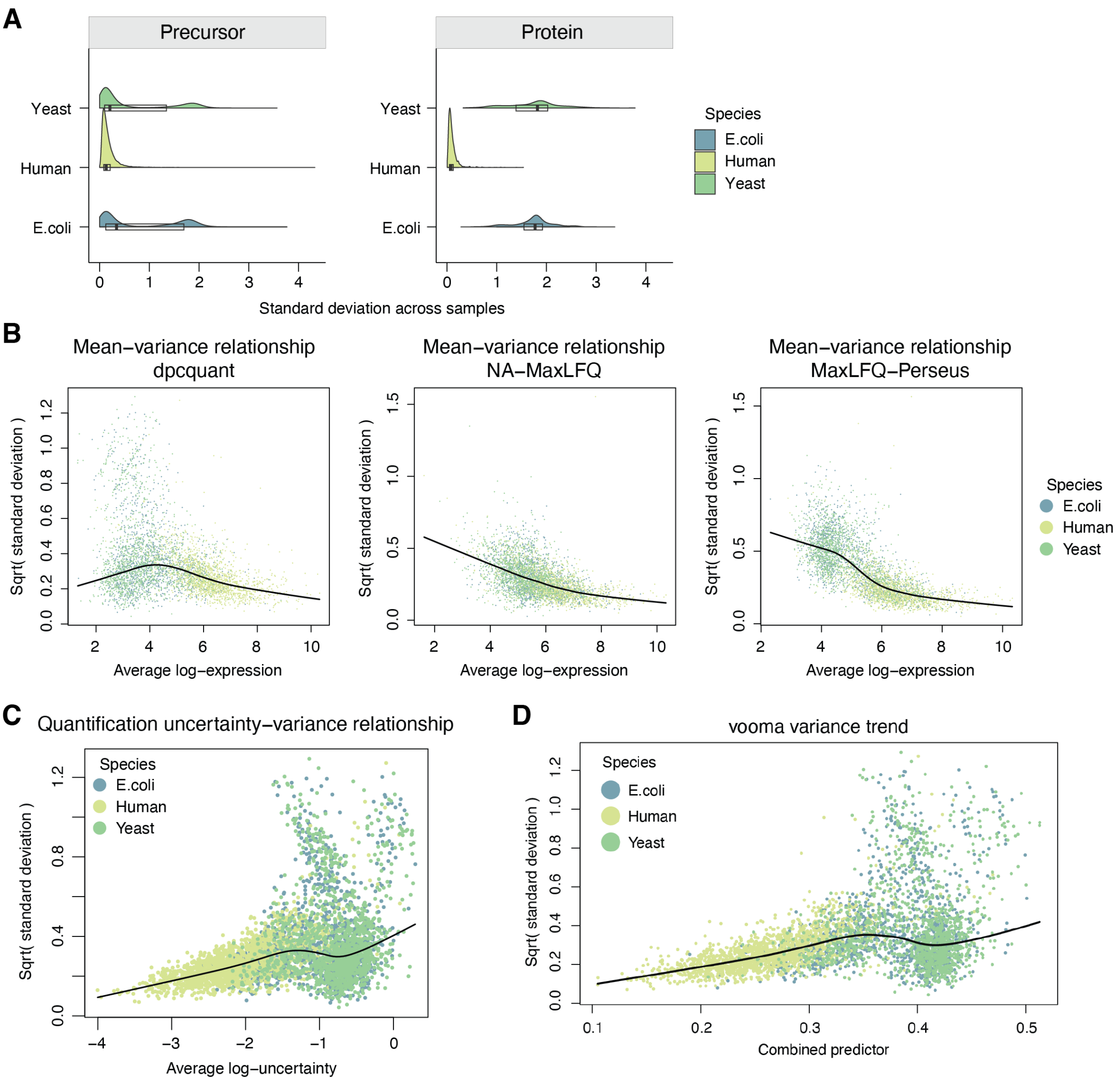


**Figure S13:** For mixed-species data processed by DIA-NN, (A) precursor- and protein-wise standard deviations by species, (B) mean-variance relationship, (C) quantification-uncertainty relationship, (D) vooma variance trend on protein quantification by DPC-Quant.
